# Nested admixture during and after the Trans-Atlantic Slave Trade on the island of São Tomé

**DOI:** 10.1101/2024.10.21.619344

**Authors:** Marta Ciccarella, Romain Laurent, Zachary A. Szpiech, Etienne Patin, Françoise Dessarps-Freichey, José Utgé, Laure Lémée, Armando Semo, Jorge Rocha, Paul Verdu

## Abstract

Human admixture history is rarely a simple process in which distinct populations, previously isolated for a long time, come into contact once to form an admixed population. In this study, we aim to reconstruct the complex admixture histories of the population of São Tomé, an island in the Gulf of Guinea that was the site of the first slave-based plantation economy, and experienced successive waves of forced and deliberate migration from Africa. We examined 2.5 million SNPs newly genotyped in 96 São Toméans and found that geography alone cannot explain the observed patterns of genetic differentiation within the island. We defined five genetic groups in São Tomé based on the hypothesis that individuals sharing the most haplotypes are more likely to share similar genetic histories. Using Identical-by-Descent and different local ancestry inference methods, we inferred shared ancestries between 70 African and European populations and each São Toméan genetic group. We identified admixture events between admixed groups that were previously isolated on the island, showing how recently admixed populations can be themselves the sources of other admixture events. This study demonstrates how complex admixture and isolation histories during and after the Transatlantic Slave-Trade shaped extant individual genetic patterns at a local scale in Africa.

## Introduction

The Trans-Atlantic Slave Trade (TAST) was one of the largest human migrations of historical times, involving the forced displacement of more than 12 million enslaved Africans according to historical sources [1]. Population genetic studies have extensively investigated the impact of these forced migrations on the genetic diversity of admixed populations on either sides of the Atlantic [2–6]. These studies provided important contributions for the understanding of genetic admixture dynamics in the context of the TAST and European colonization, as well as for the general understanding of human genetic admixture processes [7–9].

Admixture histories are rarely simple, with distinct populations coming into contact only once over time. Numerous admixed populations descended from enslaved Africans experienced several periods of recurring introgressions from populations originating from different regions of Africa [3,8,10]. Moreover, often in the colonial context, the enslaved and non-enslaved communities were reproductively isolated and subject to strict demographic constraints. Over time, changes in the economic viability of plantations, shifts in colonial policies, and the abolition of slavery contributed to evolving socio-cultural contexts, resulting in new forms of community integration and segregation [11]. The aim of this study is to account for successive admixture and isolation histories on the island of São Tomé, in light of the documented history of settlement and social segregation during five centuries of Portuguese colonization.

The archipelago of São Tomé e Príncipe, in the Gulf of Guinea, was uninhabited at the time Portuguese sailors arrived on the island of São Tomé around 1470 and rapidly settled the island from 1493 onwards [12]. On this island, the colonizers established for the first time an economic system based on plantations and the exploitation of enslaved labor for the massive production of sugar cane, known as the Plantation Economic System, that was later deployed throughout European colonies in the Caribbeans and the Americas [11]. Furthermore, the archipelago served as an important hub for the Trans-Atlantic Slave Trade between the 16th century and the abolition of the TAST and slavery during the 19^th^ century [13]. Historical records show that enslaved Africans were initially traded from the Niger delta in the kingdom of Benin, in a region that is now Nigeria. However, as the local demand for enslaved labor-force increased during the 16^th^ century, together with the growing trade with colonies from the other side of the Atlantic, São Toméan merchants expanded their slave recruitment areas southwards, to the kingdom of Kongo and other regions of northern Angola [14].

In the 19^th^ centuries, São Tomé e Príncipe became an important producer of coffee and cacao. Following the abolition of slavery in 1875, the plantation-based economy relied this time on the exploitation of indentured servitude. Contractual workers, known as “*serviçais*”, were mostly recruited from other Portuguese colonies, including Angola, Mozambique and Cabo Verde. Remarkably, over the course of the 20th century, approximately 80,000 individuals from the archipelago of Cabo Verde in West-Western Africa migrated to São Tomé [15]. As São Tomé e Príncipe, the Cabo Verde archipelago was also colonized by the Portuguese crown in the 1460’s and underwent more than 400 years of history linked with the TAST [16]. Nevertheless, the regions of origins of enslaved Africans and the histories of colonial exploitation and plantation economy differed significantly between the two archipelagos [17].

The island of São Tomé provides unique opportunities to disentangle the impact of social segregation on genetic diversity in the colonial context of the TAST, throughout the expansion of the Plantation Economy system, and after the abolition of the TAST and that of slavery. Extant communities within São Tomé are associated with different histories related to marooning, slavery, and post-slavery migrations, which are also reflected in the linguistic landscape on the island. Since the beginning of Portuguese colonization, the São Tomé forested interior of the island served as a refuge for runaway slaves, who remained largely isolated in maroon communities until the 19^th^ century, when one of these communities became known as the *Angolares* [18]. Today, *Angolares* communities in the north-eastern and south-western coasts of the island are known to speak Angolar, one of the two autochthonous creole languages of the island [19,20]. In parallel, freed enslaved-Africans became the largest social group in São Tomé during the 17^th^ and the 18^th^ century, a period of profound economic decline and relative abandon of the colony by Portuguese settlers and by the Portuguese crown [21]. Their descendants are often associated with the Forro creole language, “Forro” literally meaning “freed slave” [20]. Finally, descendants of Cabo Verdean *serviçais* may be considered to form a third historical community associated with yet another creole language spoken on the island: the Cabo Verdean Kriolu [22].

In this complex context of forced and deliberate migrations to São Tomé and segregation within the island, previous genetic studies have identified substantial genetic structure with a limited number of loci [23,24] or samples [5]. Coelho et al. suggested that the identified genetic structure within São Tomé was mainly caused by a strong signal of genetic differentiation between *Angolares* and non-*Angolares* individuals, with increased genetic drift having occurred in the former group. More recently, Almeida et al. generated exome sequence data in 25 individuals from the islands of São Tomé and Príncipe, and found similar genetic contributions from both the Gulf of Guinea and Angola in individuals speaking the Angolar and Forro creoles in São Tomé. The nested genetic structures and the mosaic of African genetic diversity found by these previous studies in São Tomé thus raised a series of fundamental questions about the histories of admixture in the island. What are the origins of the genetic ancestors of São Toméans? Are the diverse genetic ancestries evenly distributed among present-day São Toméans? Within São Tomé, and spanning existing linguistic-anthropological communities, when and how did genetic admixture and isolation processes occur?

In this study, we investigated 96 unrelated individuals sampled in 13 sampling sites from São Tomé, each genotyped at 2.5 million SNPs genome-wide. We identified five genetic groups of individuals in the sample based on haplotype-sharing patterns. We inferred the shared-haplotypic ancestries for each five São Toméan genetic groups using previously published genome-wide data from 70 African and European populations, including an extensive sample from the archipelago of Cabo Verde [6,25–28]. We reconstructed recent shared ancestries between admixed groups using sharing patterns of long tracts Identical-by-Descent. We finally propose a chronology of successive admixture and isolation events that may explain the genetic diversity observed in São Tomé, using model-based ancestry inference approaches. This study highlights the influence of the complex history of European colonization, including TAST, social segregation, and post-slavery migrations, on extant genetic patterns in São Tomé.

## Results

### São Tomé genetic diversity in the global context

To investigate the genetic diversity across the island of São Tomé in the worldwide context, we combined our genotype data on 96 unrelated individuals from São Tomé (Fig 1) with a reference dataset of population samples from Africa, Europe, and the Americas, including other populations descended from enslaved Africans in Cabo Verde, African-Americans in southwest USA (ASW) and African-Caribbeans in Barbados (ACB) (S1 Table) [6,25–28].

**Fig 1.**
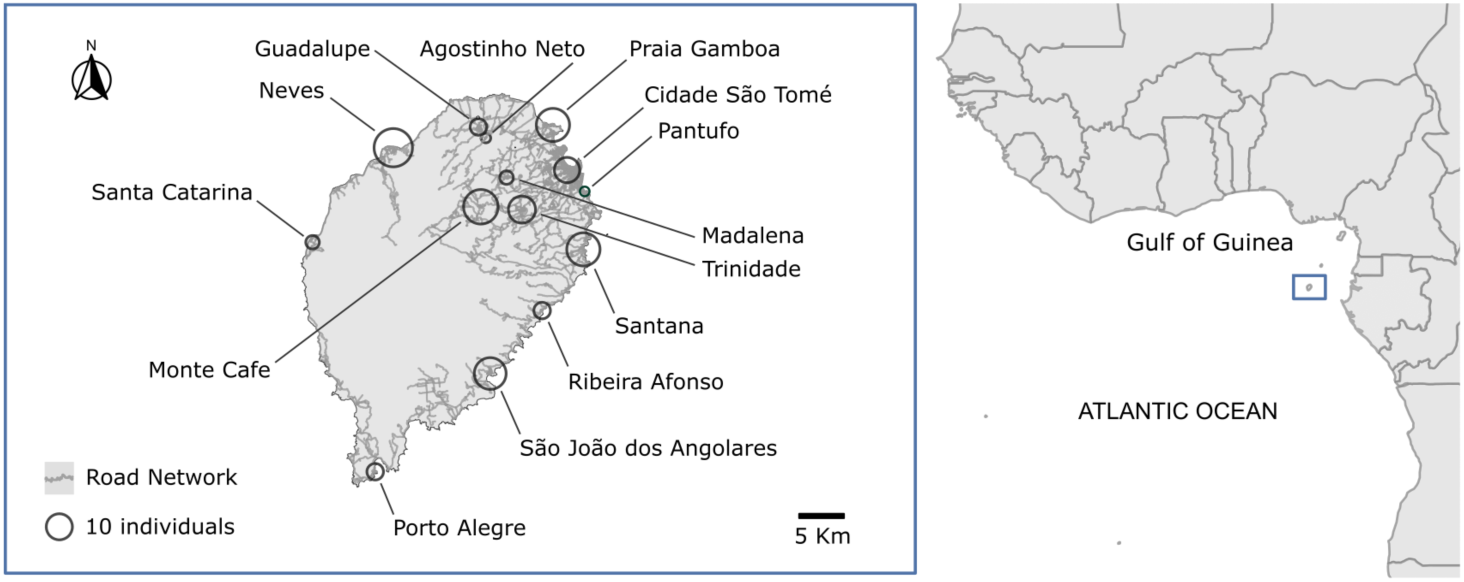
The island of São Tomé in the Gulf of Guinea. Distribution of individual samples collected from 13 locations across the island of São Tomé. The road network of the island is represented in grey. Each circle represents a location, with the size of the circle proportional to the sample size. On the right, the map shows the position of São Tomé in the Gulf of Guinea.

Fig 2A shows the sampling locations of the 70 populations included in the reference dataset, which are grouped into 10 distinct geographical regions. We consider a subset of population samples from previous studies, focusing particularly on regions in Africa that were used as key ports of embarkation during the Trans-Atlantic Slave Trade [1] (S4 Table). Fig 2B shows the first two dimensions of a Multi-Dimensional Scaling analysis calculated on Allele-Sharing Dissimilarities (ASD) [29], between each pair of individuals in our data set including 411,121 autosomal SNPs. The first axis of variation is explained by major genetic differentiation between African and European populations, while the second axis captures genetic differentiation within Africa, revealing a north-south gradient ranging from West-Western African to South African populations.

**Fig 2.**
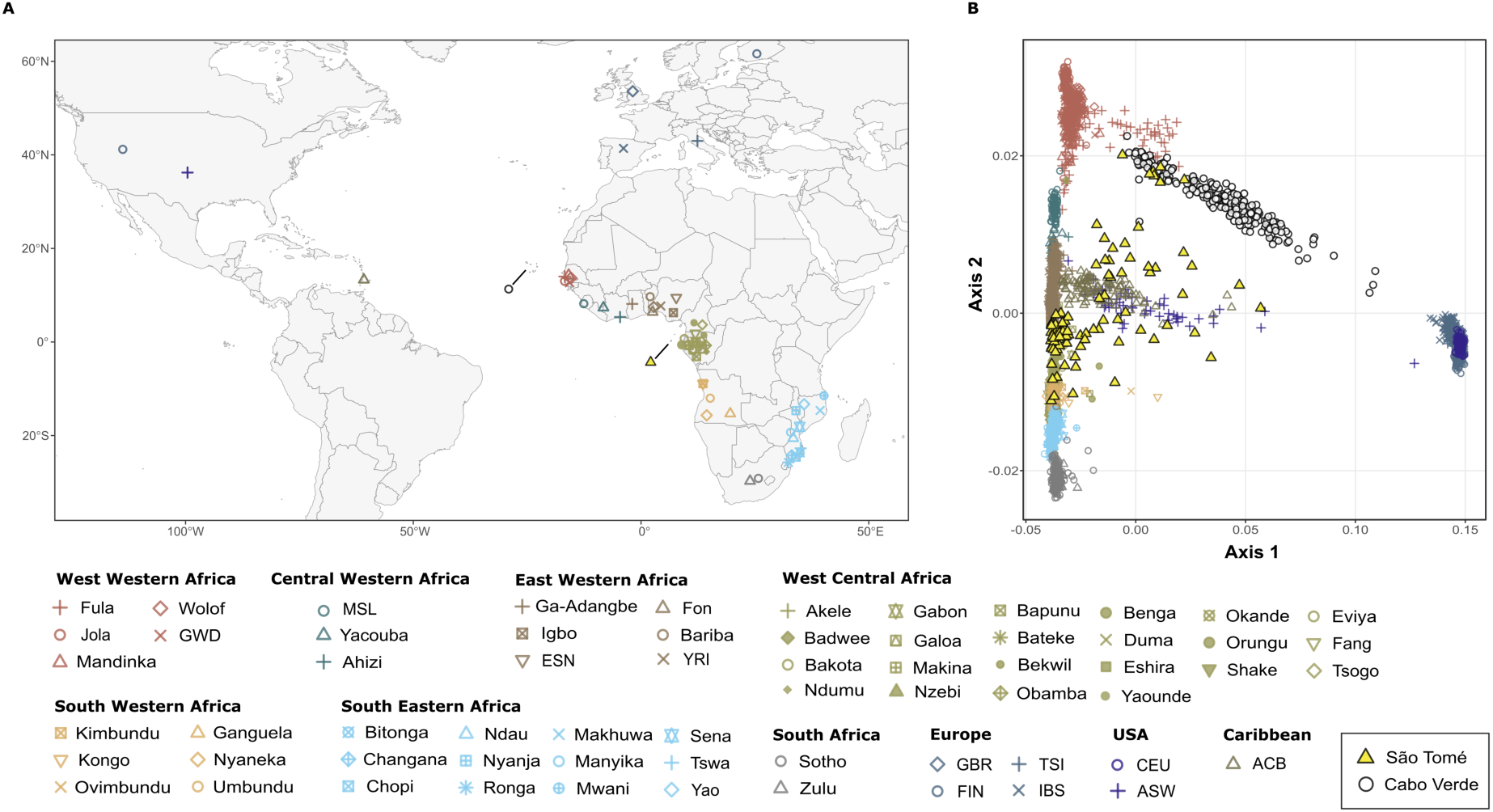
The genetic differentiation of the worldwide population samples. **(A)** Geographical locations of the 70 population samples considered in the present study. **(B)** MDS projection of pairwise allele sharing dissimilarities (ASD) among São Toméans, Cabo Verdeans and other African, American and European populations. The first two axes of variation in the MDS are calculated among 3203 individuals using 411,121 autosomal SNPs. On the bottom, the color-coded legend sorts the populations into 10 distinct geographical regions in continental Africa, Europe and the Americas.

The 96 unrelated individuals born in São Tomé are highly dispersed across the two first dimensions of the ASD-MDS, compared to the 70 population samples from Africa, Europe, and the Americas (Fig 2B). Other populations descended from enslaved Africans show a relatively simpler pattern, in spite of being also highly admixed: Cabo Verdean individuals fall along a trajectory going from European to West-Western African populations, reflecting the impact of slave recruitment from the neighbor areas of Senegambia and Guinea-Bissau in their current genetic profile [6,30]; African-Americans (ASW) and African-Barbadians (ACB) cluster along a trajectory going from Central-Western African to European populations, in agreement with a different history of recruitment of slaves mainly in the Gulf of Guinea [6,31].

We further applied an unsupervised clustering algorithm, ADMIXTURE [32], to visualize several axes of inter-individual genetic variation and population structure (Fig 3). The results of the ADMIXTURE analysis reflect the patterns of genetic diversity observed in the ASD-MDS, while providing further information on genetic differentiation within Cabo Verde and São Tomé at higher values of *K*.

**Fig 3.**
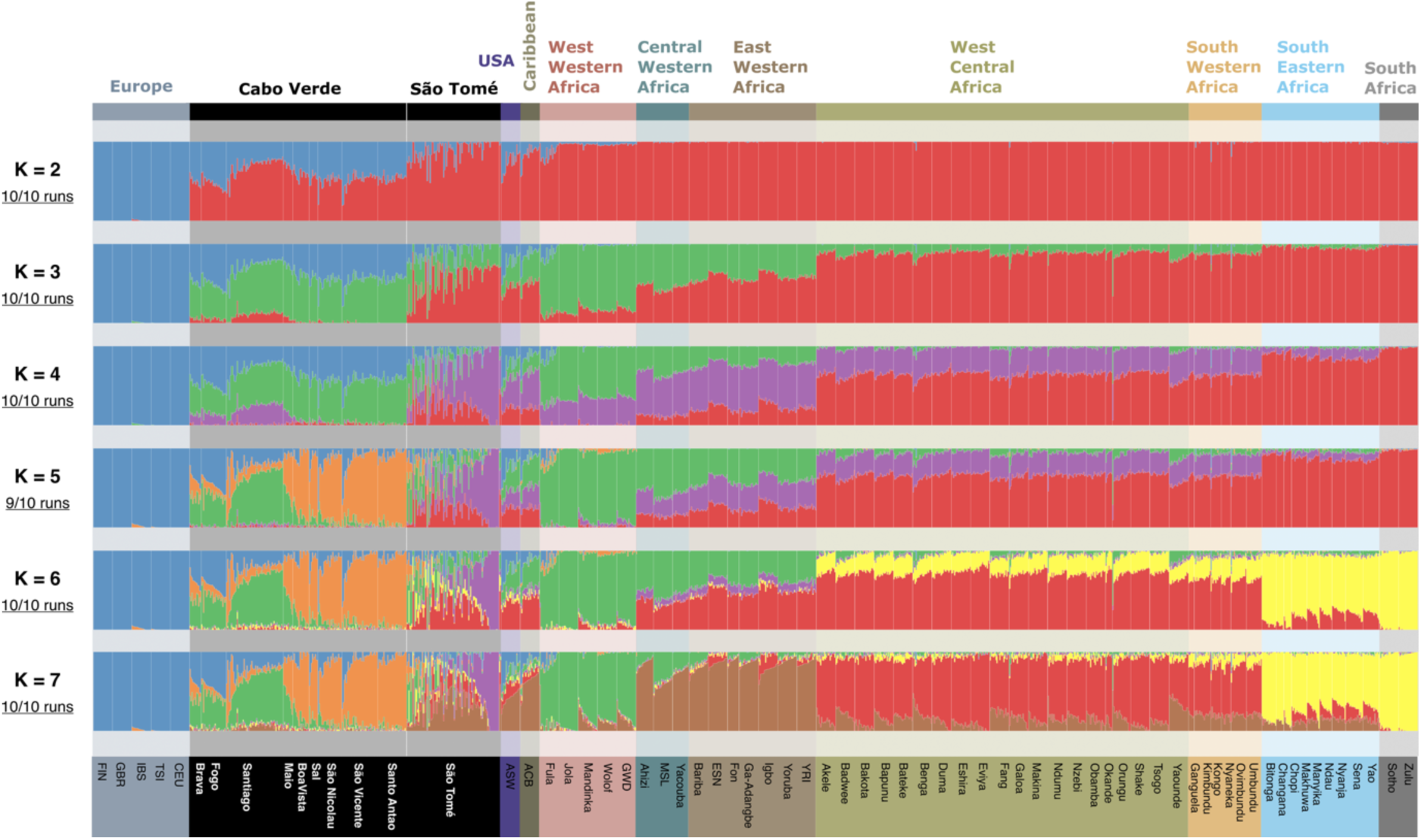
Unsupervised ADMIXTURE analysis. Unsupervised ADMIXTURE analysis using 110,499 LD-pruned (*r²*<0.1) autosomal SNPs from 1347 unrelated individuals, including 96 individuals that reported to be born in São Tomé, 2 individuals born in Príncipe, 225 individuals born on seven different islands in Cabo Verde, and a random sample of 20 individuals for each of the remaining population samples (S4 Table). The proportion of independent ADMIXTURE runs closely resembling one-another is indicated on the left of each panel below each *K* value. Population samples are ordered by geographical regions according to the color code indicated in Fig 2.

As for the first dimension of MDS, ADMIXTURE clustering at *K*=2 is explained by differences between African and European populations, maximizing the red and blue clusters, respectively. At higher values of *K*, new clusters capture differences among African populations. At *K*=7, West-Western African populations maximize the green cluster, and East Western African populations maximize the brown cluster. Also, at this value of *K*, West Bantu-speaking populations from West Central and South Western Africa present the highest proportions of the red cluster, while East Bantu-speaking groups from South Eastern and South Africa maximize the yellow cluster.

In general, genetic clusters at *K*=7 discriminate among different areas of slave recruitment in Africa, revealing genetic similarities between populations currently living in these areas and populations descended from enslaved Africans. However, while African-Barbadians (ACB) and African-Americans (ASW) display relatively simple patterns with resemblance mainly to Central and East Western Africa, Cabo Verde and São Tomé exhibit more complex genetic structures that cannot be simply explained by the diverse African origins of their enslaved settlers.

In Cabo Verde, while the southern islands of Fogo, Santiago, and Brava show high proportions of the green cluster found in West Western Africa, individuals from the northern islands (Santo Antão, São Vicente and São Nicolau) maximize the orange cluster, which is found predominantly there, and might have resulted from *in situ* differentiation due to high genetic drift [6,30,33].

In São Tomé, some individual profiles are close to that of Cabo Verdeans, while others present varying proportions of clusters found in East-Western, West-Central and South-Western Africans. Finally, another group of individuals has a unique genetic pattern, represented by the violet cluster, which is not observed outside the island.

### Haplotype-based population structure within São Tomé

We analyzed patterns of shared haplotypes between all sampled individuals from São Tomé using ChromoPainter2 and fineSTRUCTURE [34]. By looking at the patterns of the number and length of shared haplotypes, we identified five different clusters of individuals (C1-C5) with high levels of inter-individual haplotypic resemblance (Fig 4).

**Fig 4.**
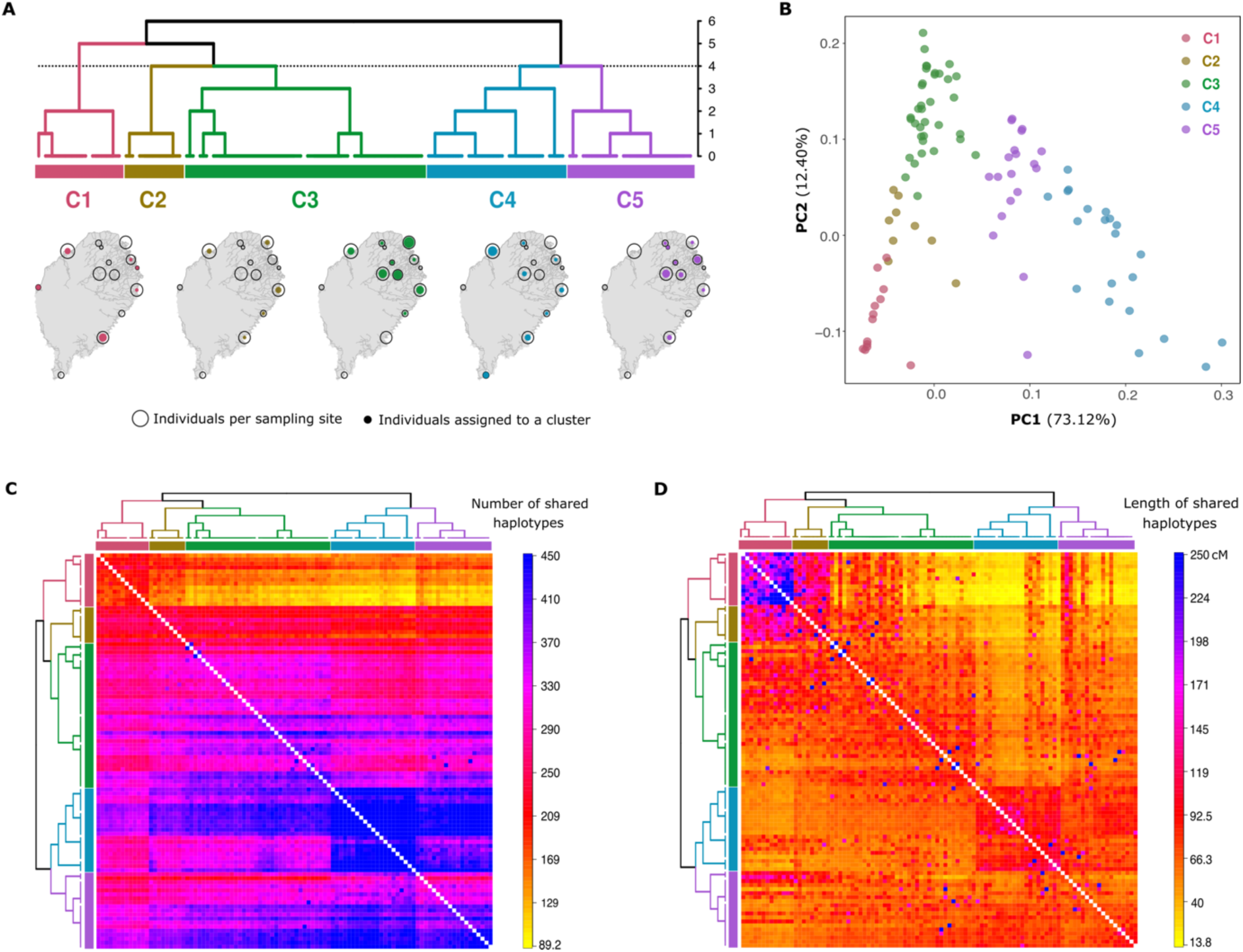
Genetic clustering and population structure in São Tomé. **(A)** Clustering of 96 individuals from São Tomé into five genetic clusters using the fineSTRUCTURE algorithm based on the number of haplotype segments shared between each pair of individuals. The vertical axis of the dendrogram represents the distance or dissimilarity between clusters. The geographical distribution of individuals within each cluster is illustrated below the dendrogram. The size of empty black circles indicates the number of individuals sampled at each location, while the size of colored dots represents the number of individuals per cluster at each location. **(B)** Principal Component Analysis (PCA) based on the co-ancestry matrix represented in panel (C). Individual labels are color-coded according to the genetic clusters identified in the fineSTRUCTURE dendrogram in Panel (A). **(C)** Co-ancestry matrix based on the total number of haplotype chunks that each individual (row) copies from any other individual (column). **(D)** Co-ancestry matrix based on the cumulated length (in cM) of haplotype segments that each individual (row) copies from any other individual (column).

Fig 4A presents a dendrogram displaying the phylogenetic relationships between the five clusters, as well as their spatial distribution across São Tomé. The majority of individuals sampled from the same locations are found to belong to more than one genetic cluster. This indicates that the geographical location of sampled individuals does not fully account for the observed genetic clustering patterns. Instead, it suggests that these patterns may be influenced by variation in admixture proportions. In a PCA calculated on the same co-ancestry matrix used to infer the fineSTRUCTURE dendrogram, individuals grouped in clusters C1 and C4 are separated along the first PC, while the second PC separates individuals belonging to the C3 cluster (Fig 4B). Within this triangle, C2 individuals lay in between C1 and C3 along the second PC, and C5 individuals lay in between C4 and the rest of São Tomé individuals along the first PC.

Moreover, we found that C1 individuals mostly share haplotypes with each other - both in terms of total number (Fig 4C) and cumulative length (Fig 4D) - and copy relatively few haplotypes from other individuals on the island. On the contrary, C2 individuals copy relatively long haplotypes from C1 individuals, as well as several haplotypes from the rest of São Toméan individuals with varying cumulative lengths. Individuals of the C3 cluster copy relatively more haplotypes from themselves than from the rest of the island, while individuals from clusters C4 and C5 differ markedly from other clusters, sharing the highest number of haplotypes among one another.

### Haplotypic ancestry of genetic clusters

The haplotypic ancestry of each São Toméan cluster was modelled as a mixture of the possible source populations included in the reference dataset using the Chromopainter2-SOURCEFIND pipeline [35].

Based on the co-ancestry matrix computed with Chromopainter2, we inferred a maximum of eight source populations per “Target” population sample using SOURCEFIND. First, we considered each one of the admixed populations descended from enslaved-Africans as a separate “Target” population, and all other populations from continental Africa and Europe as “Donors” (Fig 5A). In agreement with previous studies [3,4,6,27,30], we found that the genetic composition of island populations from Cabo Verde essentially resulted from admixture between Europeans (from 27% in Santiago to 51% in Fogo) and African individuals from West-Western Africa (from 46% in Fogo to 69% in Santiago), while African-Barbadians (ACB), and African-Americans (ASW) resulted from admixture between Europeans (15% in ACB, 24% in ASW) and Africans originating in Central and East-Western Africa (49% in ASW, 82% in ACB), South-Western Africa (3% in ACB, 18% in ASW), and West-Western Africa (9% in ASW only). São Tomé as a whole displays a more complex ancestry profile, including contributions from Europe (9%), West-Western Africa (16%), East-Western Africa (37%), South-Western Africa (36%), as well as South-Eastern Africa (2%). Importantly, when Cabo Verdean populations are considered as possible “Donors” (Fig 5B), the haplotypic ancestry from the island of Santiago in Cabo Verde (15%) replaces the Mandinka GWD haplotypic ancestry and part of the Iberian IBS component that were identified previously in São Tomé (Fig 5A).

**Fig 5.**
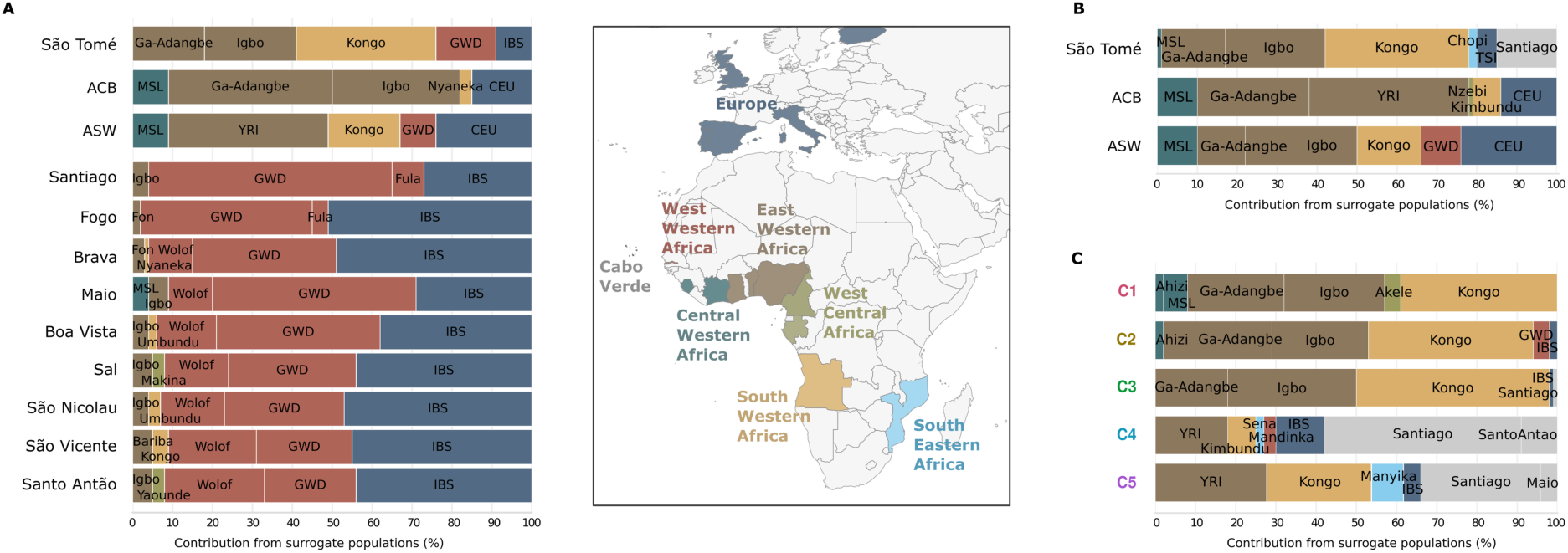
Haplotypic ancestry of São Toméan and other admixed populations descended from enslaved-Africans. **(A)** SOURCEFIND results showing the shared haplotypic ancestry between each São Toméan (STP), Barbadian (ACB), Afro-American (ASW), and Cabo Verdean population samples, set as Target, and 57 populations from continental Africa and Europe, set as Donors. **(B)** SOURCEFIND results showing the shared haplotypic ancestry between each STP, ACB, and ASW population samples, set as Target, with 66 populations from Africa and Europe, including Cabo Verdean populations from nine islands, set as Donors. **(C)** SOURCEFIND results showing the shared haplotypic ancestry between each São Toméan cluster, set as Target, with 66 populations from Africa and Europe, including Cabo Verdean populations from nine islands, set as Donors. The Donor populations are color-coded by geographic regions in Africa and Europe, as indicated in the legend at the center of the figure.

In Fig 5C, each of the five São Toméan genetic clusters (C1 to C5), is considered as a separate “Target” population and Cabo Verdean individuals from seven different islands are also included as seven separate “Donor” populations. Interestingly, clusters corresponding to the most basal bifurcation of the fineSTRUCTURE dendrogram (C1-C3 vs. C4-C5, Fig 4A) have different ancestry profiles: the C1, C2 and C3 clusters share haplotypic ancestries with Kongo-speakers from Angola, in South-Western Africa (from 39% in C1 to 48% in C3), and the Ga-Adangbe from Ghana and the Igbo from Nigeria, in East-Western Africa (from 49% in C1 to 51% in C2); the C4 and C5 clusters share haplotypic ancestry with Cabo Verdeans, mostly from the island of Santiago (30% in C5, 49% in C4). Interestingly, we found evidence for small amounts of shared haplotypic ancestry between the C4 and C5 cluster and the Sena and Manyika population from Mozambique, in South-Eastern Africa (2% and 8%, respectively).

Taken together, these observations suggests that the major division between the São Toméan clusters (C1-C3 *vs*. C4-C5) is due to variable contributions from different source populations, while further splits occurring within each major division may be related to subsequent episodes of *in situ* isolation and gene flow.

### Long IBD-tracts shared among São Toméans and Cabo Verdeans

We explored the sharing of identical by descent (IBD) tracts of the five genetic clusters in São Tomé with each other, and with each of the island populations of Cabo Verde. Fig 6A summarizes sharing patterns of IBD tracts longer than 18 cM, representing approximately the last eight generations of recombination between individuals [2]. We observe differences in IBD sharing patterns between and within each São Toméan group. Notably, individuals of the C1 cluster share the highest total length of long-IBD tracts between each other, and with individuals of the C3 cluster. Individuals of the C2 cluster share relatively low levels of long IBD tracts with each other, while sharing long IBD tracts with both the C1 and C3 clusters. Individuals of the C4 and C5 clusters are the only ones to share long IBD tracts with Cabo Verdean populations, in accordance with the SOURCEFIND results (Fig 5), although the C5 cluster shows higher values of total length of long IBD tracts shared with the C1 and C3 clusters in São Tomé, when compared to the C4 cluster.

**Fig 6.**
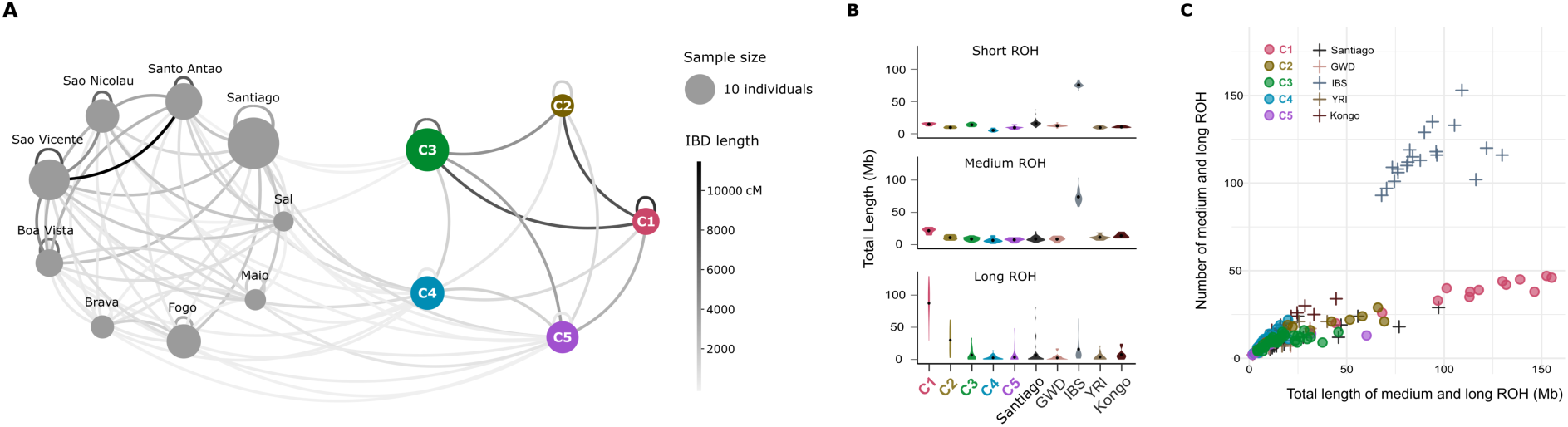
IBD tracts sharing and ROH analysis. **(A)** Network displaying the cumulative length of Identical by Descent (IBD) tracts longer than 18 cM shared within and between population samples from São Tomé (right) and Cabo Verde (left). The size of the points is proportional to the size of the population sample. The color of the lines is proportional to the total cumulative length of long IBD tracts (in Mb). **(B)** Violin plots illustrating the total length of Runs of Homozygosity (ROH) within each cluster of São Tomé, the Santiago population in Cabo Verde, and other African and European populations. **(C)** Scatter plot where each point represents an individual sample, depicting the number of medium and long ROH plotted against their total length in Megabases (Mb).

### Runs of Homozygosity

We further investigated the presence of Runs of Homozygosity (ROH), arising when an individual inherits IBD segments from a recent common ancestor. ROH of different lengths are usually generated by different demographic processes: medium ROH can be generated by recent demographic events such as bottlenecks, while long ROH are usually due to recent parental relatedness [36–38]. Fig 6B reports the total length of long, medium and short ROH for each population separately, and Fig 6C reports the total length of long and medium ROH versus their total number for all populations, calculated with GARLIC [37]. C1 individuals present a total length of long ROH higher than any other population considered. The relationship between the number and size of ROH highlights that overall C1 has few, very long ROH, suggesting that it underwent inbreeding [39]. The sum of total length of long ROH in C2 individuals is intermediate between the C1 and C3 clusters in São Tomé. The C4 and C5 clusters present similar average levels of medium and long ROH as the Santiago population in Cabo Verde.

### Dating admixture events for each São Toméan cluster separately

We inferred the timings of major admixture events that occurred in each of the five São Toméan genetic clusters, based on recombination distances between ancestry tracts using the algorithm implemented in fastGLOBETROTTER [40]. In this approach, we specified a separate set of “Surrogate” populations for each São Toméan genetic cluster, which were identified by SOURCEFIND as the “Donor” populations most likely involved in the admixture events of each cluster (Fig 5C). The fastGLOBETROTTER algorithm utilizes a mixture model for the “Surrogate” populations, allowing it to infer the relative contributions of each “Surrogate” population to two alternative mixtures, representing the first and second admixing sources for each admixture event. Overall, fastGLOBETROTTER identifies strong signals of admixture in all the five São Toméan clusters and attempts to date admixture events by considering a simplified scenario that assumes one major pulse of admixture, an important caveat that need to be kept in mind for the interpretation of the following results (See Materials and Methods and Discussion).

For the C1, C2, and C3 clusters, the major admixture event primarily involves populations from East-Western Africa, specifically the Igbo and Ga-Adangbe, on one side (Fig 7A), and the Kongo population from South-Western Africa on the other (Fig 7B). The dates of admixture events, considering a single pulse of admixture, range from 9 to 11 generations ago, and their relative Confidence Intervals are largely overlapping (Fig 7C). For the C3 cluster, the single pulse of admixture inferred 11 generations before present would correspond to the year 1694 (CI: 1636-1769) when considering a generation time of 30 years. Moreover, admixture is estimated to approximately 9 generations before present for the C1 cluster, corresponding to the year 1763 (CI: 1654-1824), and similarly for the C2 cluster, around 1748 (CI: 1683-1864). The C4 and C5 clusters likely result from varying levels of admixture between Cabo Verdeans and São Toméans. In both clusters, the contribution from admixed São Toméans is represented as a mixture of East-Western Africans and South-Western Africans (Fig 7A), and admixture events are inferred at approximately 9 generations before present, around 1760 (CI: 1736-1795) for the C4 cluster, and 1751 (CI: 1714-1790) for the C5 cluster (Fig 7C).

**Fig 7.**
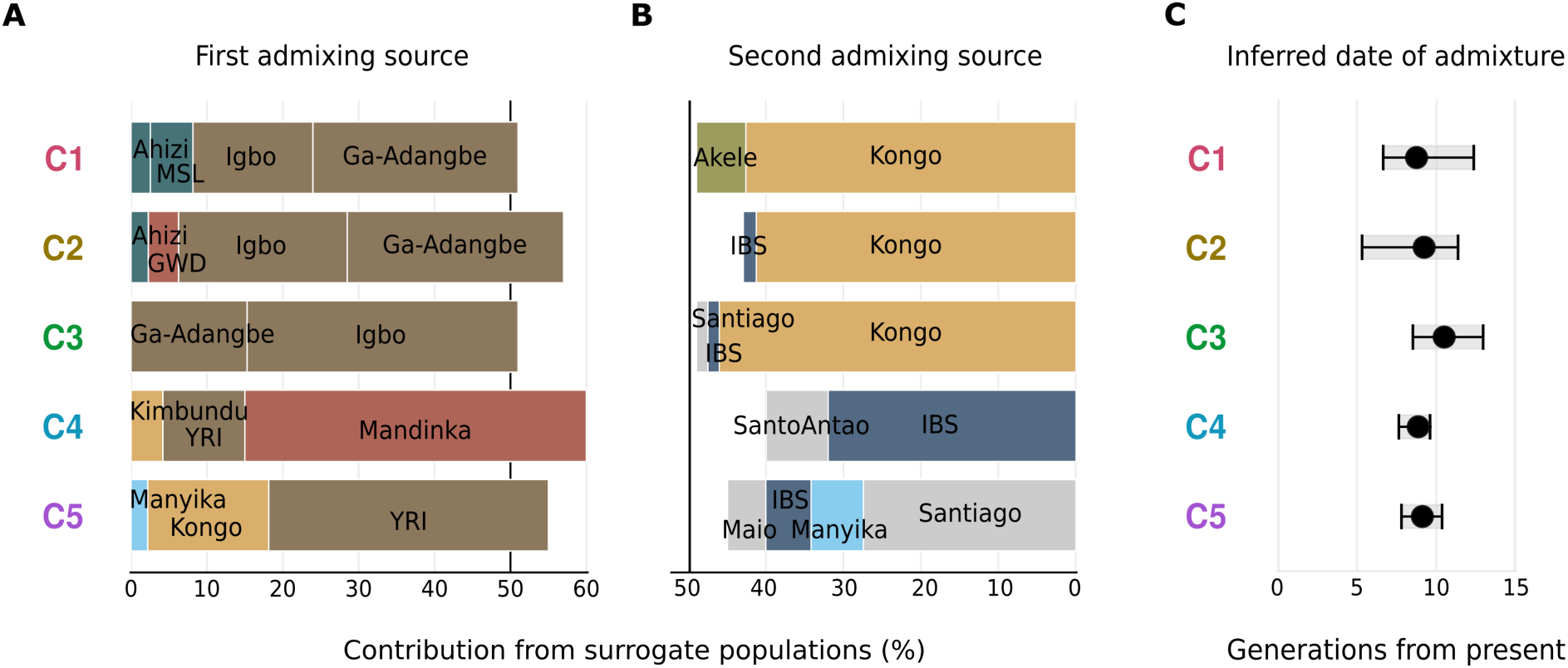
Admixing sources and timing of admixture events. Results of fastGLOBETROTTER analysis reporting the inferred composition of each admixing source for the most supported event, when assuming only a single date of admixture, for each São Toméan cluster. The surrogate populations are the ones inferred by SOURCEFIND in Fig 5C. For each São Toméan cluster, the relative contribution of surrogate populations to (A) the first and (B) the second admixing sources is depicted. The total contribution of both admixing sources sums to 100%. (C) Admixture event dates are reported in generations from present, with grey bars indicating the 95% Confidence Interval calculated with 100 bootstraps.

## Discussion

In this work, we analyzed the detailed structure of the genome-wide diversity of São Tomé - the earliest European colony in Equatorial Africa, which became a model for the Plantation Economic system that was later deployed in numerous other European colonies across the Americas [11]. Our investigation reveals that the genetic structure of São Tomé is more complex than that of several other populations descended from enslaved Africans, due to an intricate pattern of admixture and isolation events during and after the TAST.

We identified five genetic groups based on the hypothesis that individuals sharing the most haplotypes across the island may result from similar genetic histories. Notably, individuals from the same sampling site on São Tomé were often assigned to more than one genetic group, showing how social segregation in the colonial context operated at a micro-scale on the island.

A major division separates groups sharing similar proportions of haplotypic ancestries with East Western and South Western African populations (C1-C3), from groups with a clear contribution from Cabo Verde (C4 and C5) (Fig 4, 5). This differentiation is in line with the historically known recruitment of enslaved Africans from the Gulf of Guinea and Congo-Angola for the peopling of São Tomé from the 16^th^ to the 18^th^ centuries, which was followed by the arrival of Cabo Verdean immigrants for indentured labor in the 20^th^ century.

In addition to differences in the African origins of their genetic ancestors, the present gene pool of São Tomé’s inhabitants was also shaped by demographic events occurring within the island. In Fig 8, we propose an interpretation of the demographic history of São Tomé based on different types of genetic evidence, which explores the role of external population sources as well as *in situ* admixture and isolation events in shaping the genetic landscape of the island. In the following paragraphs, we discuss the relationships between the genetic data and the available historical and linguistic information that underlie our demographic model.

**Fig 8.**
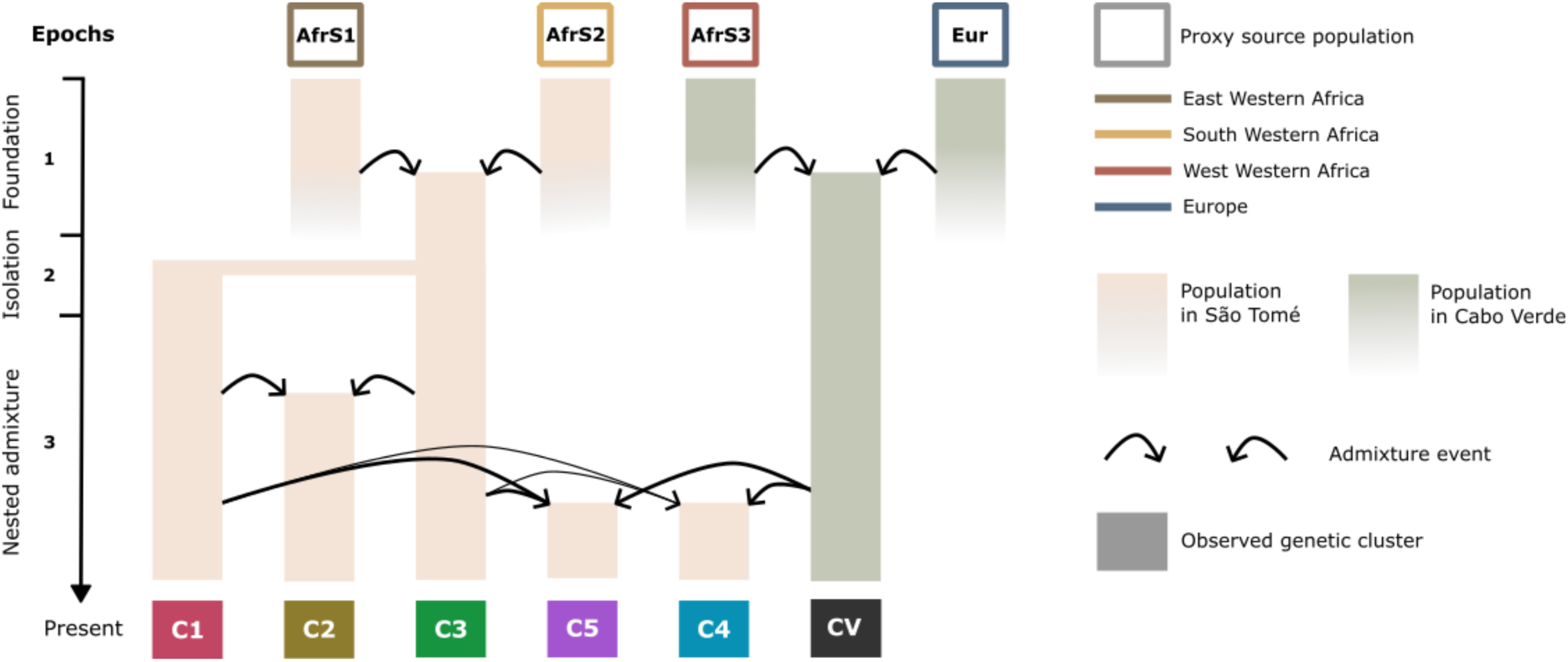
Nested admixture in São Tomé. Schematic representation of the demographic histories that may have given rise to the five genetic cluster here identified in São Tomé (Fig 4A). The timeline is divided into three epochs, spanning from the foundation of the São Toméan population to the present. Each bar represents a population evolving over time, colored according to the location of the admixture event. Proxies for admixture sources are shown at the top of the graph and are color-coded based on their origins in Africa and Europe. The colored rectangles at the bottom represents the five São Toméan genetic clusters (C1, C2, C3, C4, C5), and the Cabo Verdean population sample (CV). Each arrow illustrates the contribution of a source population at a certain epoch. In most cases, to explain the genetic diversity of a group of individuals in São Tomé, we need to consider that at least one of the source populations is itself recently admixed. We call this event “nested admixture”. Future studies should test this and similar models, including nested admixture, in their ensemble.

### A population descended from freed slaves

Towards the end of the 16th century, competition from Brazilian sugar production, coupled with significant slave revolts, led to São Tomé’s rapid economic decline and diminishing Portuguese interest in the colony [12]. Throughout the 17^th^ and 18^th^ centuries, a growing population of *Forros*, literally “freed slaves”, eventually became the dominant social group in the island [21]. Linguistically, the *Forros* speak a creole language with a Nigerian Edoid-related syntax and a predominantly Portuguese-based lexicon, which also includes various lexical items derived from Edo and Kikongo, a Bantu language from Congo-Angola [41]. Although the number of Forro creole speakers has declined in recent decades, it remains the most widely spoken creole on the island [42].

Forro-speaking individuals are currently found throughout São Tomé, with a notable concentration in the northeastern areas where the capital city of São Tomé is located [22]. Since most individuals belonging to the C3 cluster were sampled in the northeast of São Tomé (Fig 4A), it is likely that they are the descendants of the *Forros*. As noted before, individuals of the C3 cluster derive almost equal proportions of their genomic ancestry from populations currently living in Nigeria in East Western Africa, as a consequence of the slave trade in the Gulf of Guinea, and Angola in South Western Africa, reflecting the Bantu-speaking trading regions in Congo-Angola (Fig 5C). While these contributions result from extensive admixture between enslaved Africans of different origins, it is less clear when this admixture occurred.

According to the available historical and linguistic information, the features of the Forro creole that have a Nigerian origin can be traced back to the early phases of the peopling of São Tomé in the beginning of the 16^th^ century, when the slave trade was mostly limited to the Gulf of Guinea. Bantu-derived linguistic features would have entered the Forro creole in a later phase, still in the 16^th^ century, during the expansion of the plantation system, when Bantu-speaking regions became more important for slave recruitment [41]. According to our dating approach, using a single pulse model, the mixing of East Western and South Western African genetic contributions in the C3 cluster would have occurred around 1694 (CI: 1636-1769) (Fig 7C), substantially later than the mixing of linguistic features and the timing of arrival of slaves from the African mainland would suggest.

However, the dates of genetic admixture must be considered in the light of methodological limitations. In particular, fastGLOBETROTTER cannot always distinguish between discrete pulses and continuous admixture: if the period of continuous admixture is short, the model may infer a single admixture event, and the estimated date may fall within the time frame of the actual recurring admixture period [43]. Therefore, genetic admixture between East Western and South Western Africans may have occurred recurrently, both before and after the late-17^th^ century date inferred here.

### One of the first communities of runaway slaves

According to historical records, enslaved Africans escaped plantations from the onset of Portuguese colonization in São Tomé, and managed to control the mountainous interior of the island until occupation by colonial authorities in the late 19^th^ century, when one of the communities formed by runaway slaves became known as the *Angolares* [44]. Today, the descendants of the *Angolares* are mostly found in fishing communities scattered along the coast, with major settlements in the villages of São João dos Angolares and Santa Catarina where most individuals of the C1 cluster were collected (Figs 1 and 4A). It is therefore likely that the C1 cluster represents *Angolares* communities in São Tomé.

While colonial narratives have suggested that the *Angolares* descended from survivors of a shipwreck with enslaved Africans from Angola who remained almost completely isolated [44], historical, linguistic, and genetic studies have proved this hypothesis unrealistic [5,41]. The Angolar creole spoken by the *Angolares* shares many features with the Forro creole, although having a stronger lexical influence from the Bantu language Kimbundu, spoken in northern Angola. Thus, it seems likely that the two languages descend from the same proto-creole of the Gulf of Guinea and subsequently accumulated differences in the amount and type of Bantu features that became incorporated in the ancestral creole [20].

Consistent with previous genetic studies [5], we found that the *Forros* and *Angolares* of the C3 and C1 clusters, respectively, derived the most of their genetic ancestry from the Gulf of Guinea (East Western Africa, 50% and 49%) and Bantu-speaking regions in Congo-Angola (South Western Africa, 48% and 39%). However, despite this similarity, C1 individuals exhibited a markedly distinct genetic pattern, maximizing the unique ADMIXTURE violet cluster (Fig 3), and sharing very long haplotypes (Fig 4D), long IBD tracts (Fig 6A), and ROH (Fig 6B). Together, these characteristics are typical of isolated populations with small effective population sizes who experienced high levels of genetic drift and inbreeding (Ceballos et al., 2018). Similar features have also been observed in admixed population isolates from South America [45], and in other populations descended from maroon communities, such as the Noir Marron from Suriname and French Guyana [46], and another sample of *Angolares* individuals from São Tomé [5].

The strong signal of genetic drift can significantly affect recombination distances between ancestry tracts [7,47]. Nevertheless, the single pulse of admixture is estimated approximately 9 generations before present for the C1 cluster, corresponding to the year 1763 (CI: 1654-1824), which is only slightly earlier than the date inferred for the C3 cluster, and their confidence intervals largely overlap. Both clusters likely share a similar history of founding admixture, with the genetic signal of C1 more heavily impacted by subsequent genetic drift resulting from isolation and inbreeding.

It is therefore likely that the *Angolares*, sharing a major part of their genetic ancestry with the *Forros*, became differentiated through isolation after running away from the plantations (Fig 8). According to the interpretation of previous studies [5], additional contributions from runaway slaves from Angola would explain the important impact of the Kimbundu language that distinguishes the Angolar and Forro creole languages.

### Cabo Verdeans in São Tomé

The second period of Portuguese colonization in São Tomé, corresponding to the establishment of coffee and cacao plantations since the 19^th^ century, marks a profound demographic change on the island. The number of indentured laborers working on the archipelago of São Tomé e Príncipe represented approximately half of the population during the first half of the 20^th^ century [48]. According to the last census, 8% of the population in São Tomé currently speaks the Cabo Verdean Kriolu [22].

Individuals from the C4 cluster in São Tomé are more similar to Cabo Verdeans than to the rest of the São Toméan samples in terms of haplotypic diversity (S9 Fig). They share over half of their haplotypic ancestry with the islands of Santiago and Santo Antão, and present the lowest proportions of East-Western African and South-Western African contributions among all São Toméan clusters (Fig 5C). The individuals in the C4 cluster have mostly been sampled in sites close to historical *roça* plantations, such as Agostinho Neto, Monte Café, São João dos Angolares, and Porto Alegre, which hosted numerous Cabo Verdean *serviçais* to work in plantations. Moreover, they include the five São Toméan individuals that reported in our interviews that one or both their parents were born in Cabo Verde.

Our findings suggests that the Cabo Verdean *serviçais* who stayed in São Tomé after their contracts ended, and left descendants on the island, most probably came from the islands of Santiago and Santo Antão, in the south and the north of the Cabo Verde archipelago, respectively (Fig 5D). However, looking at sharing patterns of long-IBD tracts, the C4 cluster shows recent common ancestry with admixed populations from different islands in Cabo Verde (Fig 6A). Island populations of Cabo Verde share many long-IBD tracts with each other, particularly with Santiago, the first island to be settled in the 15th century, which played a key role in the founder effects that shaped the genetic diversity of the other islands [6]. SOURCEFIND potentially underestimated contributions from other islands, and likely identified Santiago as a major contributor to Cabo Verdean genetic contribution in São Tomé, partly because the algorithm was limited to using a maximum of eight surrogate populations.

The main admixture event inferred for individuals of the C5 cluster, approximately 9 generations before present (circa 1760, CI: 1736-1795, assuming 30 years per generation; Fig 7C) between West-Western African and European populations, likely reflects the founding of the admixed gene pool in Cabo Verde. However, this date is more recent than expected for the settlement of the archipelago [6].

The accuracy of admixture dating depends strongly on how well the model captures the underlying history of admixture and how well the surrogate populations represent the actual sources. The simplified admixture model used here, which assumes a single admixture pulse, does not account for subsequent, albeit minor, admixture between Cabo Verdean and São Toméan individuals. In this model, a mixture of East Western and South Western African contributions is included in the first admixture source (Fig 7A), which consists mainly of West Western African contribution, while admixed Cabo Verdean genomes contribute to the second source (Fig 7B), which consists mainly of European contribution.

### Limited European genetic contribution

Overall, São Toméan individuals exhibit limited levels of haplotypic ancestries shared with European populations, in particular compared to numerous other enslaved-African descendant populations in the Americas and in Africa [8], and even specifically compared to other populations from the former Portuguese colonial empire [3,6].

This was somewhat unexpected compared to historical records, as admixture was generally tolerated at the very beginning of the settlement as part of Portugal’s colonization efforts [49]. However, historically, the African population in São Tomé, including both enslaved and free individuals, far outnumbered the European presence on the island [50], and to a larger extent than what has been reconstructed by historians in Cabo Verde [51]. Furthermore, a substantial part of the Portuguese settlers left the island in the 18^th^ century, a period of economic crisis between the two main Plantation Economies [52].

In our haplotype-sharing analyses, individuals with European descent in São Tomé may have clustered preferentially with individuals of Cabo Verdean descent, since Cabo Verdeans have relatively high levels of European haplotypic ancestry [6]. Therefore, we know that we overestimate European haplotypic ancestry in São Tomé when we do not consider Cabo Verdeans as sources of admixture (Fig 5A), while we may underestimate it if individuals with higher European ancestry are included in genetic clusters of Cabo Verdean descent (Fig 5C). Nevertheless, it is interesting to note that the majority of European haplotypic ancestry in São Tomé actually comes from Cabo Verdean admixed genomes.

The Plantation Economic system that was originally developed in São Tomé in the 16^th^ century relied on strong social and marital segregation between free and enslaved-communities imposed by the socio-economically dominant colonists [49]. Together, these three phenomena, demography, migration, and socio-cultural constraints, likely explain the low absolute levels of shared ancestries with Europeans compared to most other populations whose history has been also intimately linked with the TAST, but where the Plantation Economy system and accompanying segregation was adopted substantially later than in São Tomé.

### Nested admixture between recently admixed populations

Beyond the three groups previously discussed—the *Forros*, descendants of freed slaves, in the C3 cluster; the *Angolares*, descendants of runaway slaves, in the C1 cluster; and the Cabo Verdean *serviçais* and their descendants, in the C4 cluster—we also identified evidence of admixture among these groups within our sample, specifically in the C2 and C5 clusters.

### Admixture between *Angolares* and *Forros*

The nine individuals of the C2 cluster are genetically closer to the C3 cluster (Fig 3, 4A), and share numerous long IBD-tracts with both the *Angolares* of the C1 cluster and the *Forros* of the C3 cluster, as well as showing intermediate levels of medium and long ROH between these two groups (Fig 6C). C2 individuals may therefore result from admixture between the *Angolares* descendants of runaway slaves and the freed population of *Forros*, after the *Angolares* communities begun to establish economic, social and cultural connections with the rest of the island population in the 19^th^ century.

Previous studies on populations descended from maroon communities emphasize strong isolation and genetic differentiation [5,46], in line with their history of escaping slavery and seeking refuge in remote, isolated locations. However, they do not account for the potential admixture with surrounding populations over time. In contrast, our findings show that marooning isolation does not necessarily imply the absence of complex admixture processes, which in turn argues for further refinement of genetic expectations for descendants of “isolated” populations in the context of admixture in general and the TAST in particular.

#### Admixture between Cabo Verdeans and São Toméans

The immigrant indentured workers were largely confined to the *roças*, or plantation units, effectively isolating them from the local São Toméan population, who refused to engage in plantation work after the abolition of slavery. This period is thus marked by the emergence of new forms of discrimination and social segregation on the island [53]. We found evidence of varying degrees of recent common ancestry between the C4 and C5 clusters, of Cabo Verdean descent, and the C3 and C1 clusters, representing the *Forros* and *Angolares,* respectively (Fig 6A). The low amount of São Toméan genetic contribution to the haplotypic ancestry of the C4 cluster aligns with the history of discrimination (Fig 5C). However, the haplotypic ancestry of the C5 cluster suggest that substantial admixture occurred between Cabo Verdeans and São Toméans (Fig 7), meaning that segregation may have been permeable or shaped by additional social factors.

#### The genetic contributions form Mozambique and Angola

In addition to Cabo Verdeans, Mozambican and Angolan contractual workers were also recruited to work on coffee and cacao plantations in São Tomé after the abolition of slavery [54,55]. While our São Toméan sample shows a substantial amount of haplotypic ancestry from Cabo Verde, we found relatively little genetic contribution from Mozambique, primarily in the C4 and C5 clusters (2% and 8%, respectively; Fig 5C). Assessing the impact of post-slavery migrations from Angola on the genetic diversity of São Tomé is more challenging, as South Western African ancestry in São Tomé may have multiple origins, including both TAST-related migrations in the 16^th^ and 17^th^ centuries, and more recent migrations of contractual workers from Angola in the late 19^th^ century.

## Conclusions and perspectives

The interplay of socio-cultural and economic factors on the island of São Tomé in the colonial context of the TAST likely influenced both reproductive isolation and gene flow between communities, with social constraints on admixture evolving over time. Our results suggest that three genetic clusters in São Tomé correspond to significant anthropological, historical, and linguistic categories - the *Angolares*, *Forros* and Cabo Verdeans - while two other clusters appear to arise from admixture among these groups. This observation reflects a gradual dismantling of a previously established genetic structure shaped by historical migrations and demographic events.

The intricate mosaic of genetic ancestries that we observed in São Tomé led us to define the concept of “nested admixture”, which refer to the gene flow between human groups that are recently admixed (Fig 8). In case the admixed source populations result from similar admixture histories, the genetic diversity of the resulting population may be composed of layers of admixture patterns, making it difficult to identify the exact source populations of the ancestry tracts. Admixture inference methods generally perform best when the source populations are not recently admixed, are genetically distinct from each other, and differ significantly from the admixed population. Interestingly, recent advances have been made to address these challenges [56]. In the case of São Tomé, future studies could attempt to infer and date admixture between the *Forros* and the *Angolares*, using, for example, the C1 and C3 clusters as proxies of source populations for the C2 cluster. Similarly, it may be possible to disentangle the two main admixture processes - the founding of the Cabo Verdean population and subsequent admixture with São Toméans - for the C5 cluster. More generally, our work highlights the need to account for gene flow between recently admixed groups, following several previous investigations on populations descended from enslaved Africans on both sides of the Atlantic [2,6,57,58].

A genealogical perspective on admixture could further illuminate on the complex patterns of genetic ancestry in São Tomé. For instance, novel theoretical developments have investigated the relationship between the number of genetic and genealogical ancestors in the Afro-American population [9,59], which would be of major interest to further understand how admixture occurred over time in the different São Toméan genetic groups here identified.

Finally, the admixture process does not occur randomly but is influenced by social and cultural factors, especially in the colonial context of the TAST [8]. Among other consequences, we may have underestimated the number of generations since admixture by assuming random mating [33,60]. Previous genetic studies suggest that in some populations descended from enslaved Africans, female and male contributions from source populations were not equal, and mating occurred preferentially between individuals with a certain genetic ancestry [33,61–63]. Future studies will need to investigate the impact of sex-biased admixture and ancestry-related assortative mating on genetic diversity patterns and admixture inference in São Tomé.

## Materials and Methods

### Genetic dataset

#### Sampling strategies

DNA sampling has been conducted by two different research groups on the two archipelagos, respectively. Sampling strategies are detailed in [6] for Cabo Verde and in [24] for São Tomé e Príncipe. Anthropological questionnaires, completed alongside DNA sampling, provide birthplace locations of each individual and their parents. DNA was collected with buccal swabs and extracted from saliva samples following standard protocols.

#### Genotyping and Quality Control

We genotyped 361 DNA samples (261 samples from Cabo Verde and 100 samples from São Tomé e Príncipe) in 5 batches, separately, using different versions of the Illumina HumanOmni2.5Million-BeadChip genotyping array read with an iScan using BeadScan software at the OMICS platform of the Institut Pasteur (S1 Table). We conducted a comprehensive genotyping quality control in four phases (S2 Table).

In Phase 1, Genotype calling, we called genotypes using Illumina GenomeStudio Genotyping Module version 1.9.4 for each batch separately. We used GenCall version 7.0.0 with a low cutoff of 0.15. We removed markers with ambiguous positions on the human genome reference sequence, and we removed markers on sex chromosomes and mitochondrial DNA. We filtered autosomal SNPs that failed the 95% call rate and 0.2 cluster separation cutoffs, and samples that failed the 95% call rate cutoff. Transversions (A/T and G/C markers) and markers with duplicated positions were also removed.

In Phase 2, Batch merging, we only kept markers genotyped in all 5 batches using *ad hoc* Python scripts. To check the concordance rate of genotype calls, we genotyped 32 individuals separately on two, three, or four different batches. We removed SNPs that were called differently on different batches for the same duplicated individual sample. We then removed the duplicated individual samples with the highest missing rate. Finally, we remapped markers positions based on rsIDs according to dbSNP Human Build 154 Release on GRCh38 genome assembly (Genome Reference Consortium Human Build 38). We removed markers whose rsIDs have multiple positions in the same build, or that do not have a position in this build. We finally retained a total of 2,104,297 autosomal markers mapped to the GRCh38 genome assembly (hg38) from 358 samples.

In Phase 3, Genetic relatedness, we removed genetically related individuals up to the 2^nd^ degree using KING version 2.2.7 [64]. In Phase 4, SNP Quality control, we subjected the remaining autosomal SNPs to quality control filters using PLINK version 1.9 [65]. We removed markers with high missing data (10%), markers that deviate from Hardy-Weinberg equilibrium (HWE), and insertion/deletion markers. We finally retained 2,104,148 curated autosomal SNPs from 330 unrelated individuals, including 233 individuals sampled in Cabo Verde and 97 individuals sampled in São Tomé e Príncipe.

#### Merging with worldwide populations

The resulting dataset of 330 family unrelated individuals from Cabo Verde and São Tomé e Príncipe was merged with 2504 samples from 26 populations worldwide included in the 1000 Genomes project Phase 3 (International Genome Sample Resource (International Genome Sample Resource IGSR) [25], with 1307 samples from 14 African populations included in the African Genome Variation Project (EGA ID EGAS00001000959) [26], with 188 samples from 15 populations in Mozambique and Angola (E-MTAB-8450) [28], and with 1366 samples from 38 sub-Saharan African populations (EGA ID EGAS00001002078) [27] (S3 Table). For the last two datasets, we conducted the same steps of genotyping Quality Controls from raw genotyping data as described before. Moreover, we checked for genetic relatedness at the end of merging each of the four datasets. We retained polymorphic markers in the merged dataset which amounted 411,121 SNPs from 5423 family unrelated individuals.

#### Population labels and geographical regions

We assigned each individual sample from Cabo Verde and São Tomé e Príncipe to an island of birth based on individual birthplaces recorded in the family anthropology questionnaires. Out of the 233 individuals sampled in Cabo Verde, 225 were born on seven of the archipelago’s islands, while seven were born outside of Cabo Verde, in particular two in São Tomé, three in Angola, one in Brazil, and one in Portugal. One individual has been excluded due to missing birthplace information. Out of 97 individuals sampled in São Tomé e Príncipe, almost all individuals were born in São Tomé, apart from one individual born in Gabon, and two individuals born in the island of Príncipe. Finally, we retained 96 individuals born in São Tomé, including the 94 individuals sampled in São Tomé and the two sampled in Cabo Verde. The dataset containing 323 individuals born in Cabo Verde and São Tomé e Príncipe will be referred to as CVSTP henceforth.

The 5423 genetically unrelated individuals from 107 populations in the final merged dataset were grouped into 16 distinct regions across the globe, including ten regions within Africa, with CVSTP identified separately, three regions in the Americas, and one region each in Europe, South Asia and East Asia.

### Population genetics descriptions

#### Allele Sharing Dissimilarity

We investigated genetic diversity patterns between each pair of individuals based on successive subsets of populations in our dataset. We first computed a matrix of pairwise allele sharing dissimilarities including all the individuals and all SNPs in the merged dataset using the ASD software (https://github.com/szpiech/asd). Then we explored three axes of variation of Multi-Dimensional Scaling projections of various subsets of this ASD pairwise-matrix using the *cmdscale* function in R [66]. In particular, we removed East Asian, South Asian, South American, Puerto Rican PUR, Mexican American MXL, Central African, East African, and four West-Central African hunter gatherers populations by subsampling the ASD matrix before conducting the MDS projection anew. Fig 2B report the first two dimensions of the ASD-MDS based on 411,121 autosomal SNPs of 3203 individuals from 77 populations, and the third dimension is presented in S1 Fig.

#### Genetic clustering

We used the software package ADMIXTURE version 1.3 [32] to explore further genetic resemblances among individuals. ADMIXTURE analysis is sensitive to sample size heterogeneities, therefore we randomly resampled without replacement 20 individuals for each population, and we removed 7 populations with less than 5 individuals, except for the island’s populations of interest in Cabo Verde and São Tomé e Principe, where we retained all individuals for this analysis. This dataset comprising 1347 individuals from 70 populations is referred to as Working Dataset henceforth (S4 Table).

Following author’s recommendations, we filtered the initial set of 411,121 autosomal SNPs for low Linkage Disequilibrium using the *--indep-pairwise* function in PLINK with a 50 SNP-window moving every 10 SNPs, and 0.1 *r*^2^ cutoff. We thus run ADMIXTURE on 110,499 LD-pruned autosomal SNPs from the 1347 individuals in the Working Dataset. We performed 10 independent runs of ADMIXTURE for values of *K* ranging from 2 to 7 (Fig 3). We used PONG [67] to define ADMIXTURE “modes” with a greedy approach for similarity threshold 0.95. The alternative ADMIXTURE mode for *K*=5 is presented in S2 Fig. We conducted an evaluation of the cross-validation error across 10 distinct runs for 15 values of *K* and found that *K*=4 yielded the lowest error (S3 Fig). ADMIXTURE results for *K*>7 are reported in S4 Fig.

### Local-ancestry inferences

#### Phasing with Shapeit4

The phasing of individual genotypes was performed using the Segmented Haplotype Estimation and Imputation tool SHAPEIT4 (version 4.2.2) [68]. We estimated haplotypes for each autosomal chromosome separately using the HapMap Phase 3 Build GRCh38 genetic recombination map [69]. We used the default SHAPEIT4 parameters for phasing autosomal data: minimum phasing window length of 2.5 Mb, and a total of 15 MCMC iterations, including 7 burn-in, 3 pruning, and 5 main iterations.

#### Chromosome painting with ChromoPainter for fineSTRUCTURE

We used the inferential algorithm implemented in ChromoPainter v2 [34] to paint each São Toméan genome as a combination of fragments received from other São Toméan individuals. We performed a first run of ChromoPainter to estimate nuisance parameters on four chromosomes using 10 EM iterations, obtaining Ne = 792.022979835471, and global mutation rate mu = 0.0020978990636338. We thus run ChromoPainter using the estimated parameters on the whole dataset. We averaged the results for all individuals by chromosome.

#### Clustering São Tomé individuals with fineSTRUCTURE

We ran fineSTRUCTURE using the co-ancestry matrix based on the number of shared chunks between each pair of individuals previously computed with ChromoPainter. fineSTRUCTURE employs a Markov Chain Monte Carlo (MCMC) sampling approach to infer the population structure. During this process, the algorithm iteratively explores the space of possible population structures. First, 100,000 burn-in steps were performed to allow the algorithm to reach a stable state. Subsequently, 100,000 further iterations were performed, retaining samples every 10,000th iteration. Following the MCMC sampling, a tree representing the inferred relationships among individuals was constructed. The best state observed during the MCMC sampling, reflecting the optimal population structure, served as the initial state for tree inference. This step provided a hierarchical representation of the population structure, offering insights into the genetic relationships among individuals within the island of São Tomé. Finally, fineSTRUCTURE classified the 96 São Toméan individuals into 17 clusters. In order to increase the interpretability of subsequent analysis, and based on the haplotype sharing patterns of the co-ancestry matrices (Fig 4C-D), we reduced the number of identified groups to five by cutting the dendrogram at height 4. For a more detailed representation of the fineSTRUCTURE dendrogram, see S5 Fig. The PCA in Fig 4B has been calculated on the co-ancestry matrix with the *pcares* function in R. The third and fourth Principal Components are reported in S6 Fig. The heatmaps in Fig 4C and Fig 4D are produced using the function provided by the authors in the FinestructureDendrogram.R script, and are capped at 450 and 250 cM, respectively, for visualization purposes. For a visual representation of how consistently pairs of individuals are grouped together across different iterations of the MCMC process, refer to the pairwise coincidence matrix in S7 Fig.

#### Chromosome painting with ChromoPainter for SOURCEFIND and fastGLOBETROTTER

We used the phased haplotype information to reconstruct the chromosomes of each individual from Africa, Europe and the Americas in the Working Dataset as a series of genomic segments inherited from a set of Donor individuals using ChromoPainter v2 [34]. As with the previous ChromoPainter run on the São Toméan sample, we first estimated nuisance parameters using 10 E-M iterations, this time on a subset of the Working Dataset, following the author’s recommendations. We randomly sampled 3 individuals per population thus subsampling 1/10th of the entire dataset, and we selected 4 chromosomes: 1, 7, 14 and 21. We averaged the estimated values across chromosomes, weighted by chromosome size, over the 10 replicate analyses. Finally, we used the a posteriori estimated nuisance parameters to run the ChromoPainter algorithm on the entire Working Dataset. We run ChromoPainter on 411,121 autosomal SNPs from 1347 individuals of the Working Dataset to prepare input files for three distinct SOURCEFIND analyses.

For the first SOURCEFIND analysis, we run ChromoPainter setting each individual as both Donor and Recipient, except for the admixed individuals of interest Cabo Verde CV, São Tomé e Príncipe STP, African-Barbadians ACB, and African-Americans ASW, which are set as Recipient only. Importantly, each Donor population consists of a maximum of 20 individuals in the Working Dataset. We obtained the averaged values of “recombination rate scaling constant” Ne = 228.3939, and “per site mutation rate” M = 0.00076. Finally, we combined painted chromosomes for each individual in each of CV, STP, ACB, and ASW populations, separately.

For the second and third SOURCEFIND analysis, and for the fastGLOBETROTTER analysis, we run ChromoPainter setting all individuals as both Donor and Recipient, except for the admixed individuals of interest STP, ACB, and ASW, which are set as Recipient only. Importantly, the nine CV populations are set as both Donor and Recipient. We obtained the averaged values of “recombination rate scaling constant” Ne = 193.2902, and “per site mutation rate” M = 0.00040. Finally, we combined painted chromosomes for each individual in each of STP, ACB, and ASW populations, separately.

Finally, we run ChromoPainter setting all individuals as both Donor and Recipient to have a symmetric matrix for PCA computation (S9 Fig). PCA was computed using the *eigen* function in R on the normalized matrix of chunks counts considering all individuals in the Working Dataset.

#### Estimating possible sources for the admixed populations using SOURCEFIND

We applied the model-based method SOURCEFIND [35] to estimate the shared haplotypic ancestry among populations based on chromosome painting. We modeled the copying vector of each “Target” admixed individual, obtained with the previous ChromoPainter analysis, as a weighted mixture of copying vectors from a set of “Surrogate” individuals. The Target and Surrogate individuals copy from the same set of Donors in the ChromoPainter analysis, in particular all the populations included in the Working Dataset except for the Target populations of interest. Since the set of Donors does not contain any Target population, this is a “regional” analysis according to the definition given in [43].

We run three separate analyses to model different hypotheses of admixture. In the first analysis (Fig 5A), we used CV, STP, ACB, and ASW populations as Targets, and all other populations in the dataset as Surrogates. In this case, the sets of Donor and Surrogate populations coincides. We aggregated results obtained for all individuals in the CV, STP, ACB, and ASW Target populations, separately.

In the second analysis (Fig 5B), we used the STP, ACB, and ASW population as Target, and all other populations in the dataset as Surrogates, including CV island populations. Indeed, we aim to estimate also this time the relative contribution of CV populations to the haplotypic ancestry of Target populations. To reduce the influence of differences in sample size among CV birth-island populations, each CV Surrogate birth-island population is composed of a random sample of 20 individuals per island, and all individuals if the island population is composed of less than 20 individuals. We aggregated results obtained for all individuals in the STP, ACB, and ASW Target populations, separately.

In the third analysis (Fig 5C), we used the five São Toméan genetic clusters as separate Targets, and we used the same set of Surrogate populations as the second analysis. Indeed, we aim to estimate the contribution of these Surrogate populations to the haplotypic ancestry of each sub-population sample in São Tomé. We aggregated results obtained for all individuals in the C1, C2, C3, C4, and C5 Target populations, separately.

We conducted 20 independent SOURCEFIND runs for each analysis. We did not allow for “selfcopying” (self.copy.ind), meaning that each Target individual is modelled as a mixture of Surrogate individuals only. We allowed for a maximum of 8 Surrogates (num.surrogates), with 4 expected number of Surrogates (exp.num.surrogates), for each MCMC iteration, and we divided each Target individuals genome in 100 slots (num.slots) with possibly different ancestry. In other words, each of these 100 slots can be claimed by any one of the maximum 8 Surrogates selected in a given MCMC iteration. We considered 400,000 MCMC iterations (num.iterations), we discarded the first 100,000 as “burn-in” (num.burnin), and we sampled an MCMC iteration every 10,000 (num.thin). Therefore, the final results consists of 30 MCMC samples following the formula M = (num.iterations - num.burnin) / num.thin. For each Target population, we retained as the final result the inferred contributions to the mixture model of the MCMC iteration with the highest posterior probability over 20 runs.

We created categories of São Toméans based solely on genetic measures. Previous studies have also used genetic criteria to cluster individuals of the reference panel into genetically-resembling groups for use as proxies of source populations in admixture models [3,35]. Instead, in our reference panel, we maintained the categories defined by the study that first described each population sample, as we believe that this approach facilitates the anthropological interpretation of genetic results. Nevertheless, we faced challenges in distinguishing between genetic contributions from populations that share much haplotypic ancestry (S8 Fig).

#### Dating admixture events with fastGLOBETROTTER

We used fastGLOBETROTTER to infer admixture using as Surrogates the populations imputed by SOURCEFIND for each Target São Toméan cluster, and dated admixture events based on the LD decay patterns among haplotypes matching any pair of Surrogates. The fastGLOBETROTTER algorithm also checks if the data fits a single rate of decay, which suggests a single date of admixture, or multiple rates of decay, which suggests multiple dates of admixture.

The same co-ancestry matrix used for the third SOURCEFIND analysis was employed, in which each individual Donor and Target genome is allowed to copy from each other. This co-ancestry matrix is utilized by fastGLOBETROTTER to infer sources, in a manner analogous to SOURCEFIND. Additionally, a separate file was prepared in which each Target genome is copying from each individual genome in the Donor population samples. This copying vector file is employed by fastGLOBETROTTER to date admixture events. I used as Donors all the populations of the reference panel, including the Cabo Verdean populations. I used as Surrogates the populations imputed by SOURCEFIND for each São Toméan cluster.

Following author’s recommendations, I first run fastGLOBETROTTER separately for each group setting “null.ind: 1” in the parameter file, inferring admixture proportions, dates, and sources over 5 iterations, and performing 100 bootstrap re-samples to generate confidence intervals around the estimated dates. In all bootstrap resamples for all Target São Toméan cluster, no date estimate was < or equal to 1, nor > or equal to 400. Therefore, the p-value for evidence of “any detectable admixture” is 0 for each Target São Toméan cluster. I thus presented in the Results the output of a second fastGLOBETROTTER run, setting “null.ind: 0”. The results of the first and second run were consistent in terms of best guess of admixture dating.

The “null individual analysis” mitigates signals in the LD decay curve due to strong genetic drift. In the “null individual” step described by [43], GLOBETROTTER constructs a “null” coancestry curve, representing the likelihood that two DNA segments, separated by a certain genetic distance and originating from different Target individuals, match a specific pair of Surrogate populations. This process aims to eliminate linkage disequilibrium (LD) decay signals in the coancestry curves that are not due to admixture, thus improving the accuracy of date estimates for admixture events. GLOBETROTTER then adjusts the coancestry curve using the “null” coancestry curve before estimating admixture dates and proportions. This step is implemented in both GLOBETROTTER and fastGLOBETROTTER. However, fastGLOBETROTTER is better able to deal with atypically long chucks inferred by ChromoPainter in drifted groups. Specifically, fastGLOBETROTTER detects if the left end of the coancestry curve is affected by long chunks, and thus removes this part of the curve prior to model fitting. Despite these corrections, the authors advise caution, as strong genetic drift can still influence the results.

According to thresholds set by the authors, all São Toméan clusters better fit one pulse of admixture, apart from the C5 cluster which fit multiple pulses. In particular, for the C5 cluster, the additional goodness-of-fit (R2) explained by adding a second date versus assuming only a single date of admixture, taking the maximum value across all inferred coancestry curves, is 0.44, which is slightly higher than the threshold of 0.35 suggested by the authors. However, the authors further suggest to visually inspect the coancestry curves, where the x-axis gives genetic distance in cM, and the y-axis gives the probability of copying from a pair of Surrogate populations at a pair of DNA segments separated by a certain cM distance. For all pairs of Surrogate populations of the C5 cluster, there is an increased probability to copy DNA segments separated by small cM distances from any pair of Surrogates. However, the 1-date curve fit the data as well as the 2-date curve starting from 2 cM distances. We thus decided to retain the 1-date results for C5, since the 2-date results account for a pulse of admixture between Cabo Verdean populations 197 generations ago, which is highly improbable since Cabo Verdean populations have been founded in the late 15^th^ century as per historical records [51].

To estimate confidence intervals for the date of admixture, we performed 100 bootstrap resampling, and we calculated the quantiles of this distribution (2.5 and 97.5 percentiles) to determine the lower and upper bounds of the confidence interval, respectively (Fig 7C).

### Recent shared ancestry tracts

#### Long identical by descent (IBD) tracts

We used hapIBD to generate shared identity-by-descent (IBD) segments from the phased data of 96 São Toméan and 225 Cabo Verdean unrelated individuals. We used the same genetic distance map used for phasing, and we employed the seed-and-extend algorithm implemented in hap-IBD with default parameters. Initially, seed segments are identified as identity-by-state (IBS) segments longer than 2 cM. While IBS segments indicate that two individuals share a genetic segment, IBD segments specifically represent shared ancestry, where the shared segment is inherited from a common ancestor. These segments are extended iteratively, considering another long IBS segment for the same haplotype pair, separated by a short non-IBS gap. Default parameters included a maximum non-IBS gap of 1,000 base pairs and a minimum extension length of 1 cM. By allowing short non-IBS gaps, the method accounts for discordant alleles that may result from genotyping errors.

We found a total of 259,624 IBD segments shared between the 323 unrelated individuals from São Tomé and Príncipe. We filtered for IBD segments larger than 18 cM, taking into account the last 8 generations of recombination according to previous calculations [2]. By summing the length of all shared IBD segments greater than 18 cM, we calculated the cumulative IBD sharing between each pair of individuals. We thus summed the cumulative length of IBD segments shared between individuals within and between each population, specifically the 5 genetic clusters in São Tomé and the 7 populations for islands of birth in Cabo Verde. We plotted the resulting matrix as a heat map, capped at 10,000 cM (S10 Fig). In Fig 6A, we built a network based on this matrix using the “networkx” package in python.

#### Runs of Hemizygosity (ROH)

We called runs of homozygosity (ROH) with GARLIC [37], considering the five genetic clusters in São Tomé as separate populations, as well as 59 individuals from the island of Santiago, in Cabo Verde, 20 Gambian GWD individuals, 20 Iberian IBS individuals, 11 Kongo individuals, and 20 Yoruba YRI individuals. For each population separately, we ran GARLIC using the weighted logarithm of the odds (wLOD) [70], with a genotyping-error rate of 0.001, and using the same genetic distance map used for phasing, window sizes ranging from 30 to 90 SNPs in increments of 10 SNPs, 100 resampling to estimate allele frequencies, and all other GARLIC parameters set to default values.

GARLIC calculates the ROH using window sizes for ROH detection and length boundaries for ROH class detection separately for each population, depending on the specific allele frequencies and distributions of ROH lengths, respectively. We calculated the sum of ROH for each class length, for each individual (Fig 6B). We also calculated the sum of the length of all medium and long ROH, and the number of medium and long ROH, for each individual (Fig 6C).

## Acknowledgements

The authors would like to thank all the São Toméan participants who contributed to this study. We also thank the “Paléogénomique et génétique moléculaire” (P2GM) platform at the Muséum National d’Histoire Naturelle, Musée de l’Homme, for their assistance in handling biological samples and generating genetic data, and the BIOMICS platform at the Institut Pasteur for carrying out the genotyping analyses.

## Ethic statement

Research sampling protocols followed the Declaration of Helsinki guidelines and informed consent was obtained from all subjects involved in the sampling in São Tomé and Cabo Verde.

The study of the São Toméan sample was undertaken with the support and permission of the Provincial Government of the Ministry of Health of the Democratic Republic of São Tomé and Príncipe, and the Provincial Government of Príncipe.

For the Cabo Verdean sample, research and ethics authorizations were provided by the Ministério da Saúde de Cabo Verde (228/DGS/11), and the French ethics committees and CNIL (Declaration n°1972648).

## Data availability

The novel genetic data presented here can be accessed via the European Genome-phenome Archive (EGA) database accession number #XX (pending) upon request to the corresponding Data Access Committee. The dataset can be shared provided that future envisioned studies comply with the informed consents provided by the participants, and in agreement with institutional ethics committee’s recommendations applying to this data.

## Supporting Information

**S1 Table. Genotyping details for the São Toméan and Cabo Verdean sample batches.** Batch codes, sampling locations, genotyping years, and the Illumina manifest used for each batch.

**S2 Table. Overview of the Genotyping Quality Control (QC) Pipeline across four phases.** Sequential steps involved in each phase of the genotyping QC, including genotype calling (Phase 1), batch merging (Phase 2), genetic relatedness filtering (Phase 3), and population genetics QC (Phase 4). Specific actions, such as removing ambiguous markers, markers on sex chromosomes, duplicates, and markers with low call rates, are noted alongside the number of markers and samples retained after each step.

**S3 Table. Population datasets included in this study.** The original publication source for each population dataset.

**S4 Table. Population datasets included in the Working Dataset.** Populations and number of individual samples included in analyses presented in this study.

**S1 Fig.**
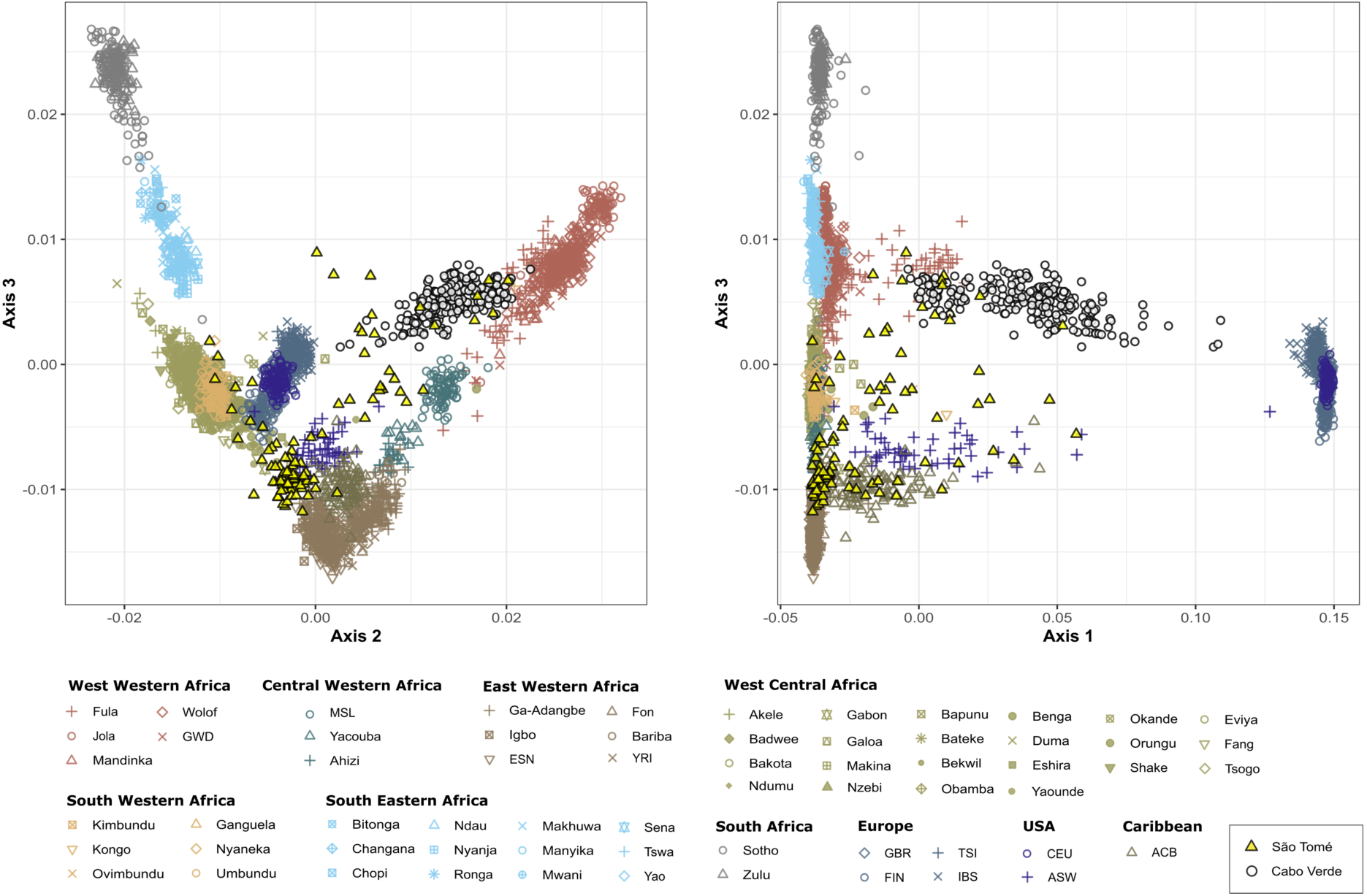
Multi-Dimensional Scaling (MDS) analysis. This figure extends Fig 2B by including the third axis of variation in the MDS projection of pairwise allele sharing dissimilarities (ASD, Bowcock et al. 1994). The MDS includes São Toméans, Cabo Verdeans, and various African, American, and European populations, with the projection based on 3203 individuals and 411,121 autosomal SNPs.

**S2 Fig.**
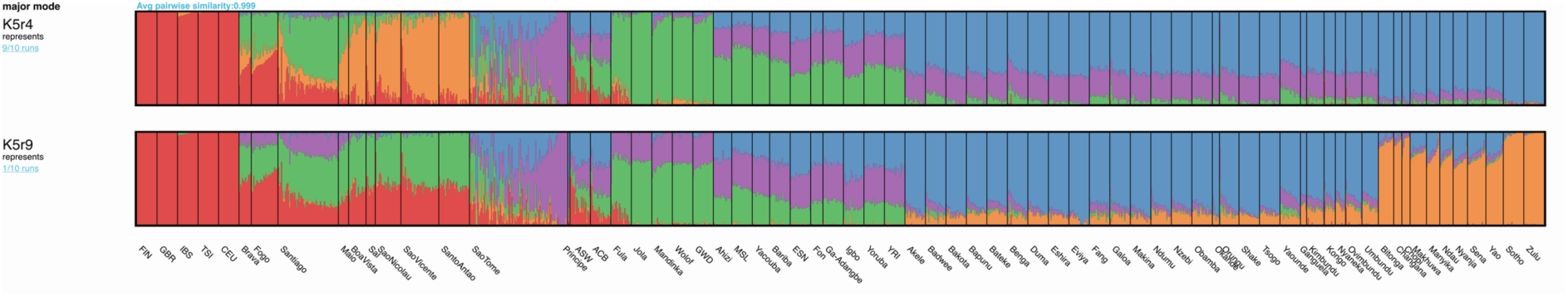
Alternative ADMIXTURE mode for *K*=5. Alternative admixture mode for *K*=5, differing from the major mode reported in Fig 3 that represents 9 out of 10 independent ADMIXTURE runs. The average similarity between this alternative mode and the major mode is 0.814581, as calculated with PONG.

**S3 Fig.**
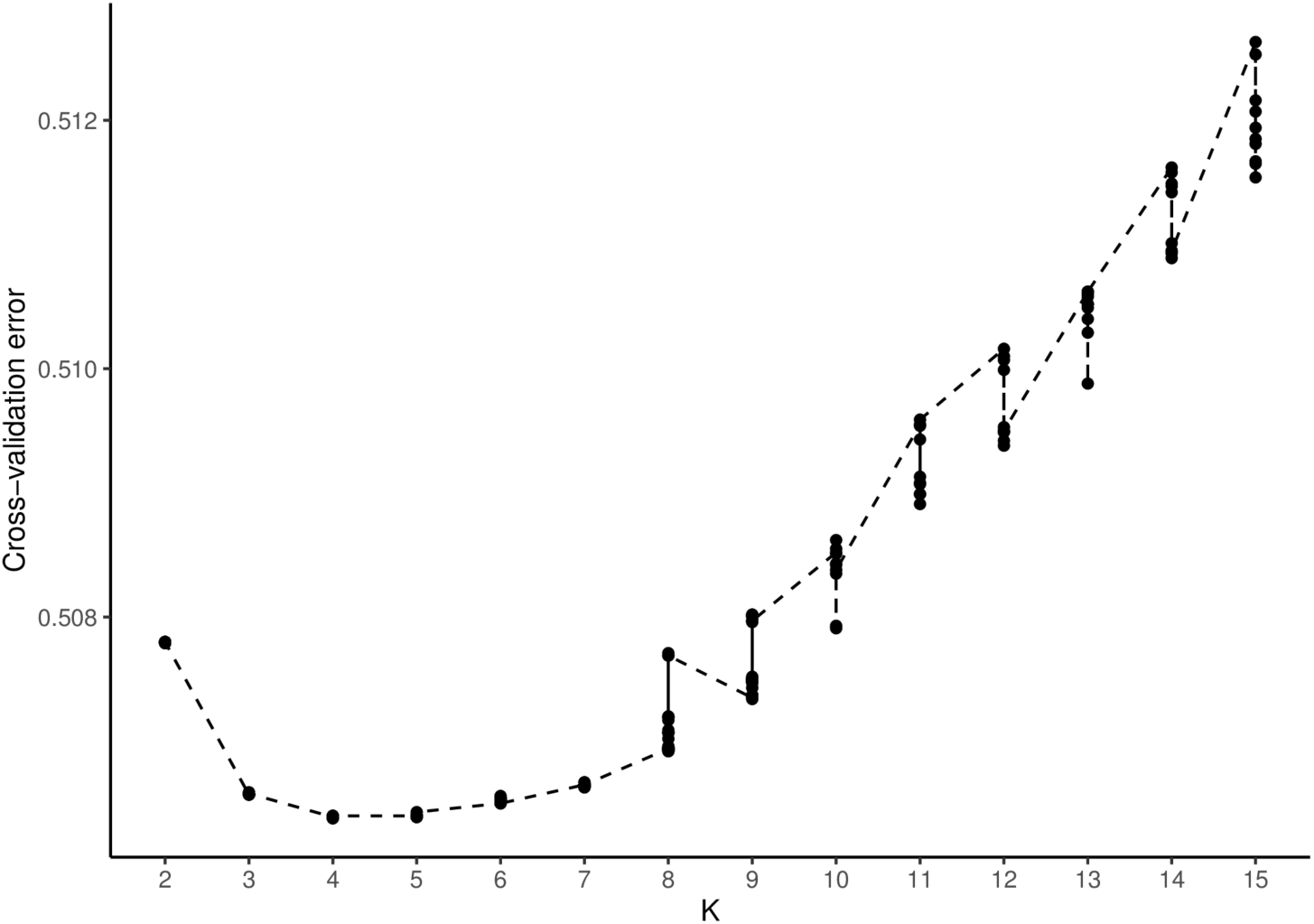
Cross-validation error of 10 independent ADMIXTURE runs for K from 2 to 15. The cross-validation error for 10 independent ADMIXTURE runs begins to increase starting from *K*=7.

**S4 Fig.**
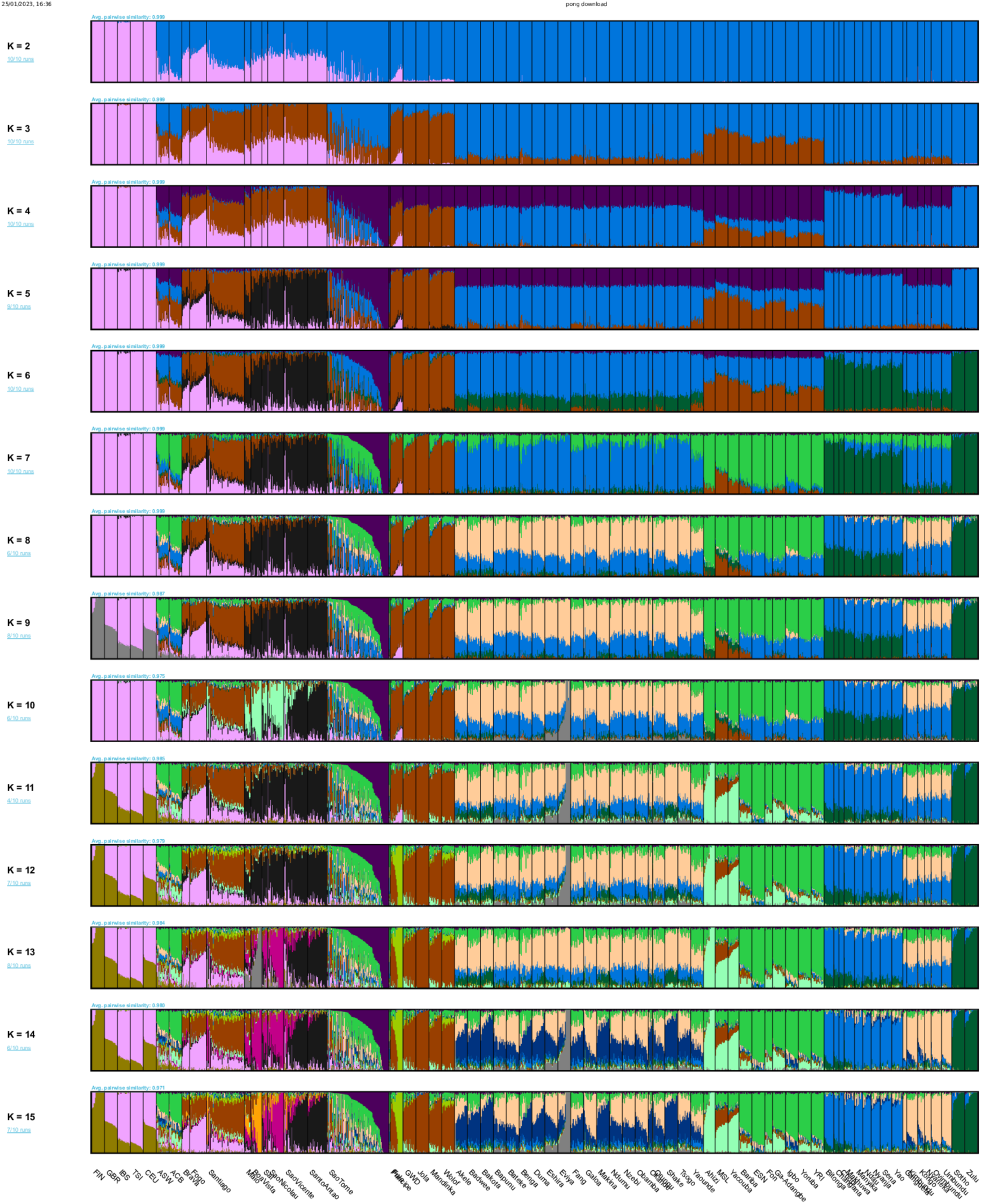
Unsupervised ADMIXTURE analysis for *K* from 2 to 15. Unsupervised ADMIXTURE analysis as in Fig 3. Here, results for *K* > 7 are reported.

**S5 Fig.**
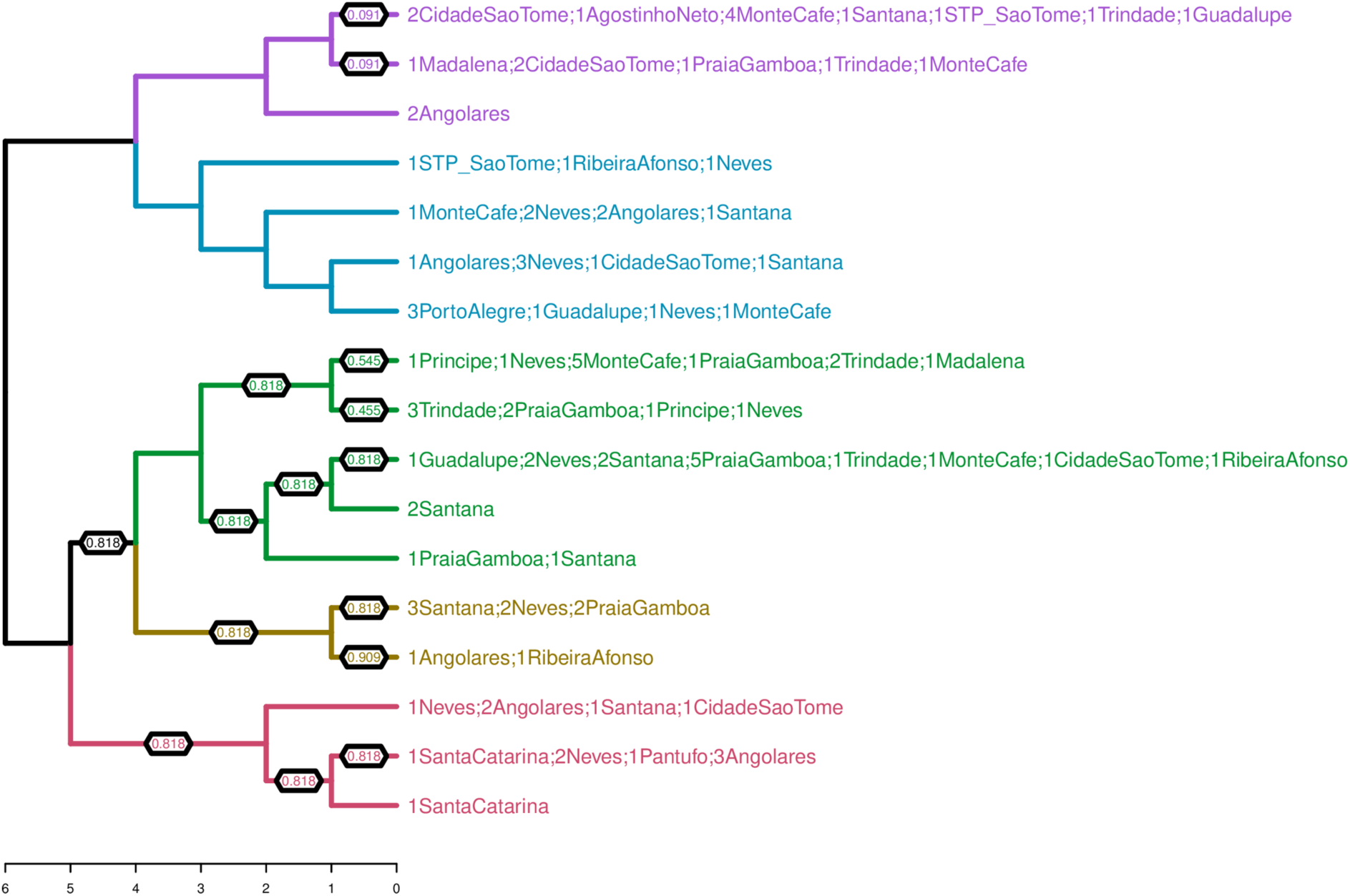
fineSTRUCTURE dendrogram of the São Toméan sample. The numbers on the edges of the dendrogram give the proportion of MCMC iterations for which each population split is observed (only displayed when the proportion is below 1).

**S6 Fig.**
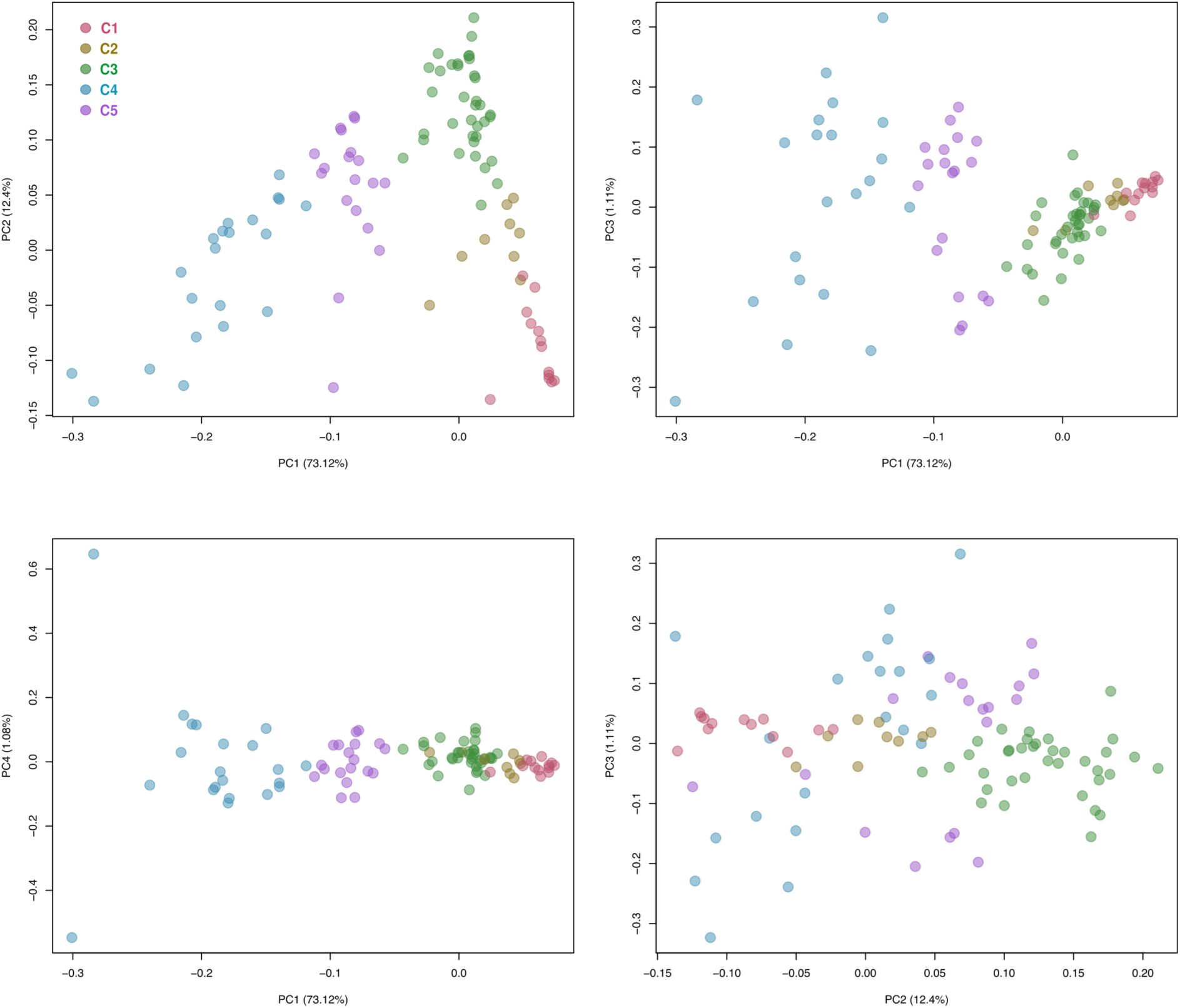
Principal Component Analysis (PCA) based on the co-ancestry matrix of the São Toméan sample. This figure expands upon Fig 4B by including additional principal components, specifically PC3 and PC4.

**S7 Fig.**
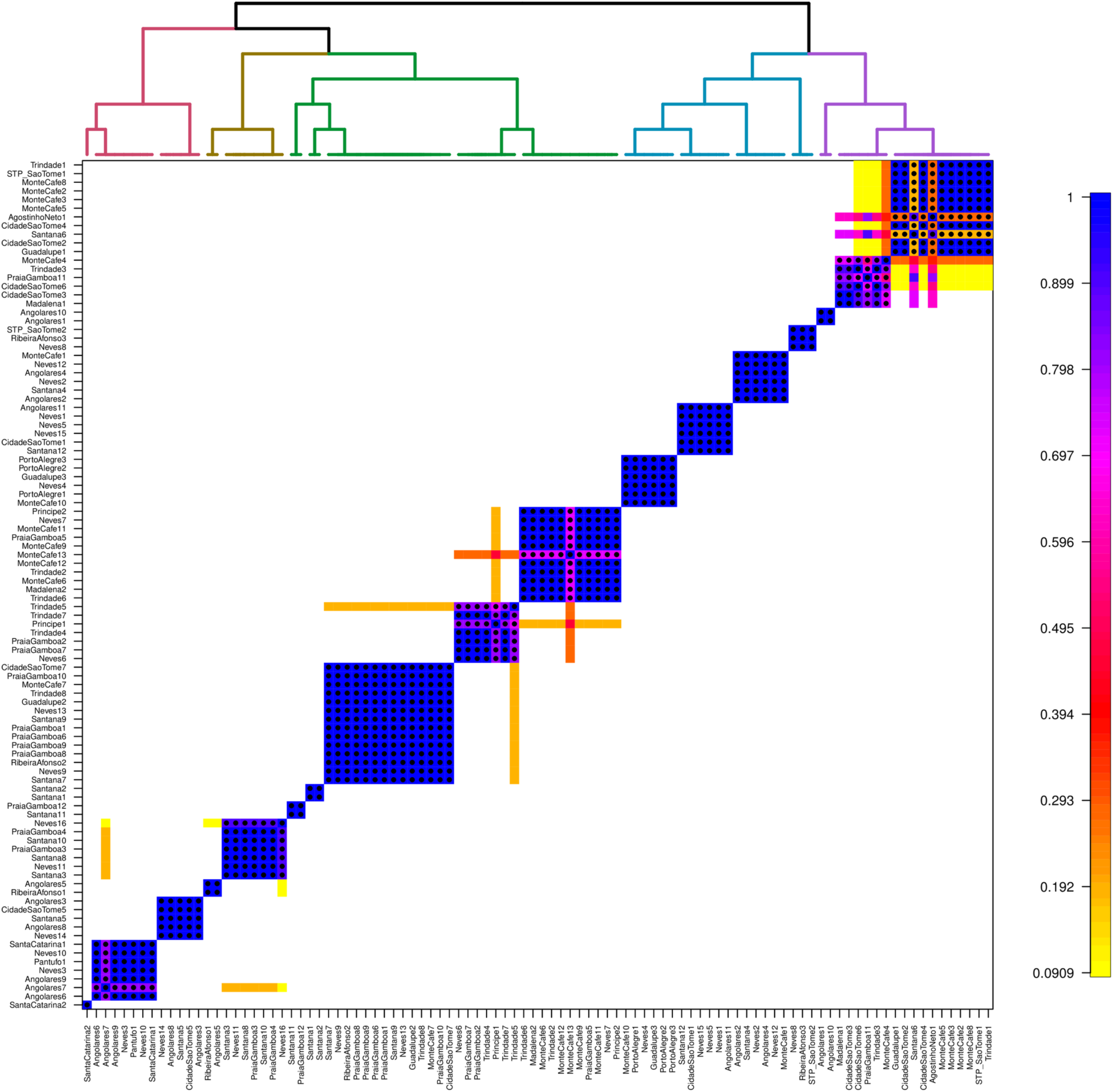
Pairwise coincidence matrix of São Toméan individual samples. The coincidence matrix is used to summarize the results of fineSTRUCTURE’s Markov Chain Monte Carlo (MCMC) clustering process. It captures how consistently pairs of individuals are grouped together across different iterations of the MCMC process. The coloring represents the average pairwise coincidence across MCMC samples. If the value is close to 1, it means that the corresponding pair of individuals is almost always grouped together in the same cluster, indicating strong genetic similarity.

**S8 Fig.**
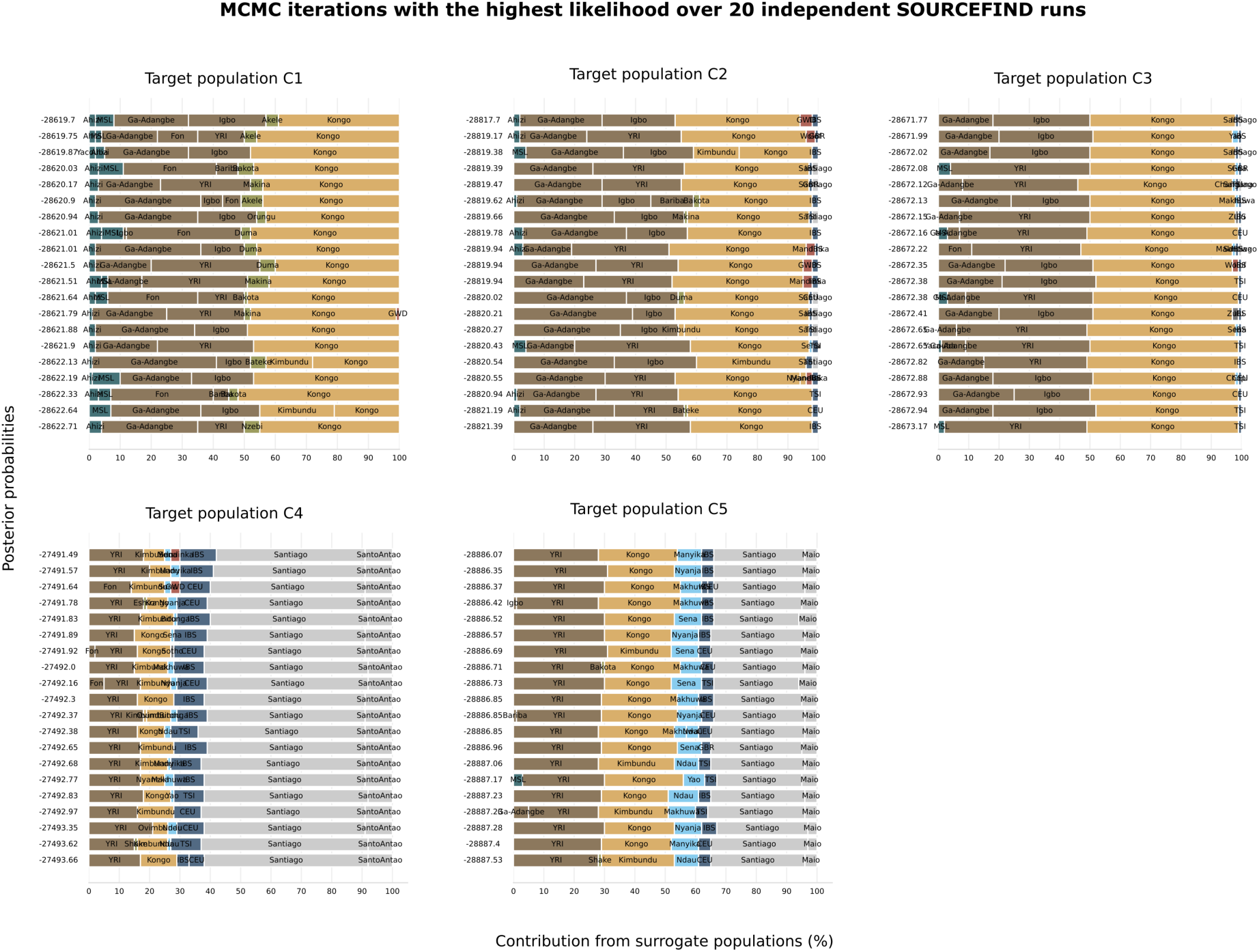
MCMC iterations with the highest likelihood over 20 independent SOURCEFIND runs. The results are ordered by posterior probability and illustrate both the consistency and variability in the source estimates across different runs.

**S9 Fig.**
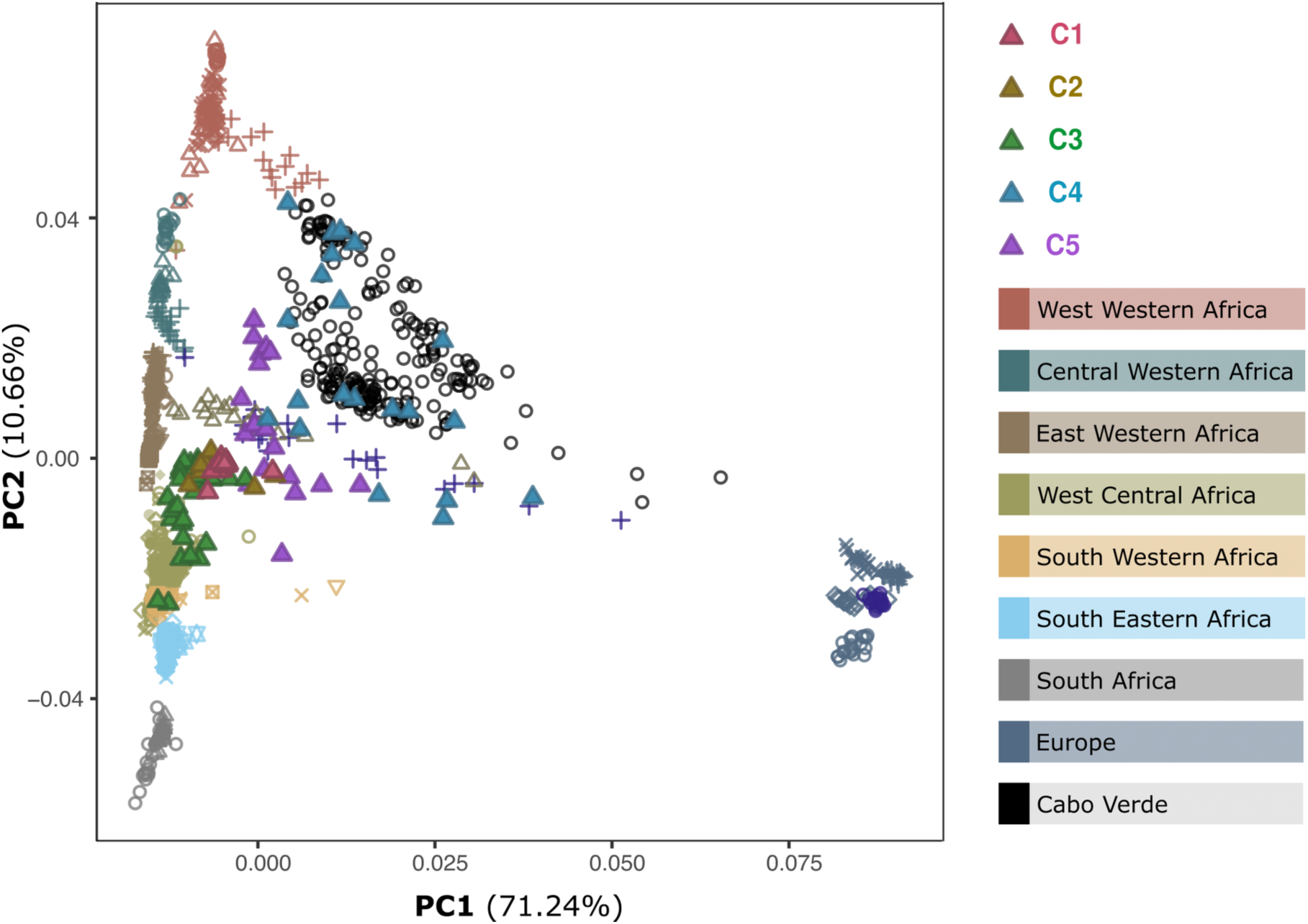
Haplotype-based PCA. Principal Component Analysis (PCA) based on chromosome painting with Chromopainter2 using all the 1347 individuals of the Working Dataset as both Donors and Recipients.

**S10 Fig.**
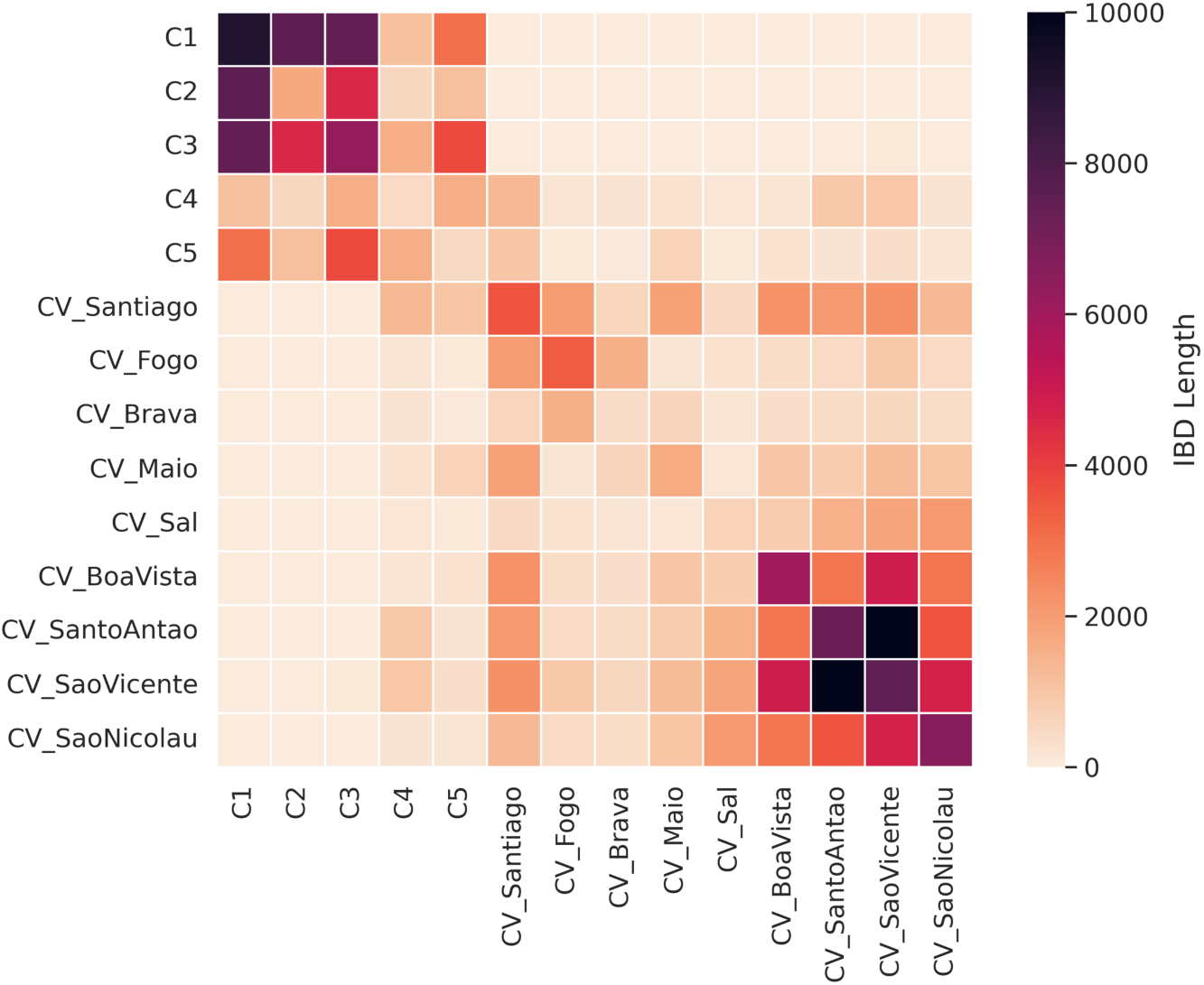
Heatmap of the cumulative length of long Identical by Descent (IBD) tracts. Alternative representation of the results shown in Fig 6A, displaying the cumulative length of IBD tracts longer than 18 cM, that are shared within and between samples from São Tomé (C1-C5) and Cabo Verde.

